# Exponential diversity-dependent diversification emerges from an individual-based model with Lotka-Volterra competition

**DOI:** 10.1101/2023.03.28.534580

**Authors:** Théo Pannetier, A. Brad Duthie, Rampal S. Etienne

## Abstract

A long-standing question in macroevolution is whether diversification is governed by the same processes that structure diversity at ecological scales, particularly competition. This competition has led to the development of a model where diversification rates depend on diversity, analogous to density-dependence in population growth models. Various versions of this model have been widely used for inference, where the rate of speciation and/or extinction can be either a linear or a power function of species number. It is, however, unknown if either approximates the diversification process that arises from the general ecological setting proposed to lead to diversity-dependence. This is of concern for inference, as failure to include a model that appropriately represents the hypothesized scenario is likely to lead to erroneous inference. Here we use an individual-based model adapted from adaptive dynamics, where fitness is governed by resource availability and the density of competitors, to determine the shape of the diversity-dependence functions. We find that the diversity-dependent rate of speciation produced by the individual-based model is best approximated by an exponential function of species diversity, consistent with a view of macroevolution where diversity increases rapidly after mass extinctions or when new adaptive space becomes available. Although we do find diversity-dependence in the extinction rate, it remains low over the entire process and erases its own signal, so it cannot be recovered from reconstructed phylogenies. The support for a linear relationship for diversity-dependent diversification found in many empirical phylogenies suggests that either our adaptive dynamics model of speciation is inadequate or there is too little information contained in reconstructed phylogenies. We indeed find evidence for the latter when pruning extinct species from our simulated phylogenies, but this does not rule out the former.

## 1 Introduction

Whether and how multi-million year evolutionary trends observed in fossil diversity and reconstructed molecular phylogenies are related to the eco-evolutionary processes taking place within contemporary communities is a challenging question that has been the focus of many studies (Raup et al. 1973; Jablonski 2008; Weber et al. 2017; Harmon et al. 2019). The model of evolution through competition and exclusion between closely related forms as described by Darwin attests that it was already a central concern in the Origin of Species (Darwin 1859; Rabosky 2013). To this day, it remains unclear how ecological interactions among organisms and their environment scale up to shape the rates of speciation and extinction that are central to macroevolutionary studies. Paleobiologists of the 1970s through 1980s proposed that the diversification of taxa within clades behaves analogously to community assembly (Raup et al. 1973; Sepkoski 1978), as described in the then-recent theory of island biogeography (MacArthur and Wilson 1967). Closely related taxa are assumed to exploit a common pool of resources available to the clade and occupy exclusive niches. If niche space is assumed to be limited, expansion of the clade results in saturation of this niche space, in turn impeding the successful establishment of new species and increasing the risk of extinction through increased competition intensity. Conversely, extinction of a taxon relaxes some of the competition pressure and provides the ecological opportunity for other species to diversify, so that in the long term, the clade tends to reach an equilibrium diversity. Under this view, diversity feeds back on diversification; i.e., diversification is diversity-dependent. The model provides a simple, intuitive explanation for how competitive interactions among individuals may have cascading effects on the formation of clades. Because it is based on the universal ecological principle of competitive exclusion, it is also applicable to any study system.

Implementations of diversity-dependence as a birth-death model have been widely used to test for a primary role of competition in a clade’s evolutionary history, both using reconstruction of fossil diversity (Silvestro et al. 2015; Ezard and Purvis 2016; Foote et al. 2018) or the branching patterns of a molecular phylogeny (Nee et al. 1992; Rabosky and Lovette 2008; Etienne et al. 2012; Weir and Mursleen 2013). Although the verbal model outlined above makes the intuitive predictions that the rate of speciation should generally decline with the number of species in the clade, and the rate of extinction should increase, it does not describe any precise relationship between the rates and the number of species, as this would require making a number of assumptions regarding the ecological setting, the nature of competitive interactions, and the mode of evolution of the species considered. Instead, it is usually assumed that the relationship between the respective per-capita rates of speciation and extinction and the number of species is simple, most often a linear function. Originating from Sepkoski (1978)’s influential study of temporal patterns of diversity in the marine fossil clades, a linear function was chosen because of tractability, and in analogy with contemporary models of island biogeography (MacArthur and Wilson 1967). Sepkoski himself acknowledged that this constituted an approximation, justified by the ease of analysis and interpretation such a model permits (for example, it is easy to predict the equilibrium diversity of the clade). Less frequently, power functions of the number of species have been used instead, often somewhat confusingly referred to as the “exponential” (Nee et al. 1992; Rabosky and Lovette 2008), or “hierarchical” (Maurer 1989; Brayard et al. 2009) diversity-dependent model. Linear and power models predict similar diversity growth curves, but differ in important aspects as a result of the curvature of both rates in the power model. First, the initial speciation rate is typically much higher in the power model, resulting in rapid (“explosive”) initial growth of the clade (Brayard et al. 2009). Second, in the power model, species diversity has a strong effect on the per capita rates when the clade contains only a few species, and a weak effect when diversity approaches equilibrium, whereas in the linear model these effects are constant throughout the diversification process (Maurer 1989; Nee et al. 1992). Finally, the dynamics of the rates beyond equilibrium diversity differ. In the linear model, speciation becomes zero, such that there exists a theoretical maximum to the possible size of the clade. By contrast, in the power model both rates are asymptotic, so that the strength of diversity-dependence itself reaches a maximum. The power diversity-dependent model has a more substantial biological background, as it was originally derived from a population-level model of “energy flow” (i.e, resource allocation), in order to establish a mechanistic foundation instead of simply assuming the shape of the relationship (Maurer 1989). These underlying population-level processes have however seldom been referred to in later uses of the model (Nee et al. 1992; Rabosky and Lovette 2008; Brayard et al. 2009).

For the purpose of statistical tests for diversity-dependence against alternative evolutionary scenarios in reconstructed molecular phylogenies, both versions of the model are often included in the set of candidate models as alternative implementations of the same hypothesis (a primary role of competition in driving diversification) (Rabosky and Lovette 2008; Weir and Mursleen 2013; Condamine et al. 2019). In this context, linear diversity-dependence is often selected over power diversity dependence. For example, Condamine et al. (2019) found that linear diversity-dependence in speciation was selected for 35 phylogenies (out of 218 phylogenies of Tetrapod families), while power diversity-dependence in speciation was only selected for 1. The choice of the form of diversity-dependence may rarely matter for the detection of diversity-dependence from branching patterns in molecular phylogenies: model selection tends to rank models with linear and power functions of diversity-dependence close to one another compared to other birth-death models (Rabosky and Lovette 2008; Condamine et al. 2019), implying that omitting either model is unlikely to change the qualitative conclusions. Yet, the selection of either model over the other leads to different interpretations of the ecological mechanisms underlying diversity-dependence (Nee et al. 1992). Importantly, other mechanisms can lead to slowdowns in the rate of diversification (Rabosky 2013; Moen and Morlon 2014), making it unclear whether what is recovered is indeed the contribution of competition, or an unrelated evolutionary scenario. Time-variable birth-death models are known to be subject to the issue of unidentifiability, where independent models representing unrelated evolutionary scenarios will have the same likelihood if they predict the same average growth of the clade (Kubo and Iwasa 1995; Louca and Pennell 2020). This issue has been recently shown to extend to diversity-dependent birth-death models (Pannetier et al. 2021). Because of this, the inclusion of several models representing the same evolutionary scenario increases the risk of inferring diversity-dependence when it is in fact absent, should the “true” scenario be congruent with either of the diversity-dependent models. This issue is not unique to macroevolution, but instead applies to model selection in general. Burnham and Anderson (2002) famously argued against automated selection procedures, and instead insisted that much attention should be given to the careful formulation of the set of candidate models. That is, the size of the set should be limited, and each model should be sufficient in itself to represent a given hypothesis. This policy has recently been advocated to address unidentifiability in molecular phylogenies (Morlon et al. 2020). Unfortunately, there is no clear justification for either form of diversity-dependence, as it remains unclear which, if any, of the models appropriately represents the effect of competition between individuals on evolutionary rates (Rabosky 2013; Weber et al. 2017). Here, we aim to address this gap, and establish what form of diversity-dependence emerges from the scenario described above. We use an evolutionary individual-based model (IBM) where differences in reproductive success depend on the profitability of a resource matched by a trait, modified by Lotka-Volterra-like competition between individuals (Pontarp et al. 2012). The model can be considered a stochastic, finite-population version of the deterministic models used in adaptive dynamics (Dieckmann and Law 1996; Geritz et al. 1998; Doebeli 2011), and such models have been used extensively to study the ecological and evolutionary conditions that lead to the formation of species (Dieckmann and Doebeli 1999; Doebeli and Dieckmann 2003). In this context, emphasis has been placed on the first branching event, and comparatively few studies have studied the macroevolutionary dynamics of such models. Yet, given enough time and depending on the model parameters, further branching may occur, such that a clade forms from an initially monomorphic population. Previous studies have described the interplay between competition and landscape dynamics on the phylogenetic structure of the resulting communities (Pontarp et al. 2012) and the shape of phylogenetic trees (Gascuel et al. 2015), and the effect of trait dimensionality on the speed of (trait) evolution (Doebeli and Ispolatov 2017). We are aware of only one study describing the relationship between the rates of speciation and extinction and the number of species in the community (Aguiĺee et al. 2018). Aguiĺee et al. provided a thorough description of the effect and interplay of biotic (competition, and the build-up of genetic differentiation) and abiotic (structure and dynamics of the landscape) factors on diversity-dependence. They found that diversity-dependence proceeded in three phases, corresponding to distinct phases of the building of communities: an initial adaptive radiation corresponding to the colonisation of all patches, followed by in-situ competition-driven diversification, and saturation of the local and global communities. Here, we consider a simpler scenario, with asexual reproduction, a 1-to-1 genotype-to-phenotype map, a unidimensional ecological trait and no spatial structure. By doing so, we seek to limit the evolutionary process to the minimal mechanisms described in the verbal model of diversity-dependence: evolutionary branching proceeds through competition-induced divergent selection and the partitioning of niche space between the resulting species, while extinction occurs as a result of stochasticity (Doebeli 2011). Total niche space and population sizes are limited, such that only a limited number of species can coexist in the final community, and diversity reaches a dynamic equilibrium (Pontarp et al. 2012). We run the simulation and measure the rates of speciation and extinction as functions of the number of species. We use the phylogenies of the resulting communities to assess whether the linear, power, or exponential function best approximates the diversity-dependent diversification process that emerges from our evolutionary individual-based model.

## 2 Methods

### 2.1 Individual-based model

We simulated evolving communities using an individual-based version of the Lotka-Volterra competition model (hereafter, LVIBM).

In this model, individuals are characterised by an ecological trait matching a resource. The fitness *W* (*z*) of an individual with trait value *z* depends on the density of competitors with a similar trait value, and on the resource abundance corresponding to this value:

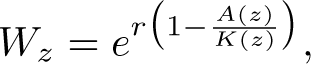

where *r* is the baseline growth rate, *A*(*z*) is the total competition intensity for an individual with trait value *z*, and *K*(*z*) is the abundance of the resource available for individuals with this trait value. *K*(*z*) is modeled as a Gaussian distribution:

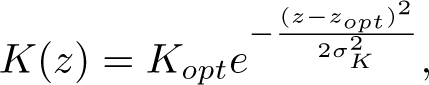

where *K_opt_* is the maximum abundance, corresponding to optimal trait value *z_opt_*, and the standard deviation *σ_K_* controls how fast the abundance of the resource declines away from *z_opt_*. The total competition intensity at trait value *z* is

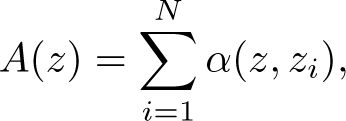

where *N* is the number of individuals in the community and,

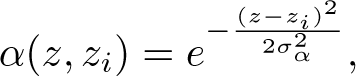

is the competition intensity experienced by a focal individual with trait value *z*, caused by individual *i* with trait value *z_i_*; 1 *≥ α*(*z, z_i_*) *≥* 0. Parameter *σ_α_* controls how fast competition between individuals declines as the trait distance between them increases.

Generations are discrete and non-overlapping, and reproduction is asexual. In each generation, the number of offspring each individual produces is drawn from a Poisson distribution with the fitness of this individual as the mean parameter. Offspring inherit the trait value of their parent, modified due to mutation; their trait value is sampled from a normal distribution with the parental trait values as mean and standard deviation *σ_µ_*,

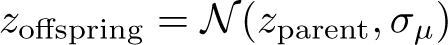

Our formulation of this Lotka-Volterra model is based on the one used by Pontarp et al. (2012), although we modified the fitness function. In Pontarp et al. (2012), the fitness function is of the form

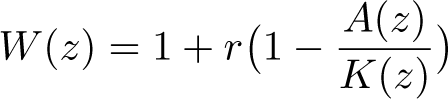

Here, we instead used the Ricker model, 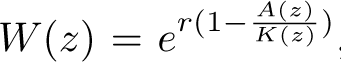, in order to avoid fitness becoming negative when 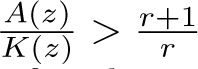, which, although unlikely and probably without much consequence for the simulation, may happen at the edges of the resource distribution. A key feature of the model is the presence of an unstable attractor in the fitness landscape, at point *z* = *z_opt_*. The phenotypes of individuals are first attracted to this point, but are eventually pushed away from it as frequency-dependence turns this fitness maximum into a local minimum. This frequency-dependence causes the population to split into two phenotypic clusters on either side of the optimum, i.e. evolutionary branching. Here, we treat such clusters as species (see next section). The new species may branch further or not, depending on *K*(*z*) and *α*(*z_i_, z_j_*), and the values of their parameters (here, *σ_α_* and *σ_K_*). Determining the conditions that lead to branching of the initial population has been the focus of many adaptive dynamics studies, and this has led to key advances in the understanding of how ecological interactions may cause speciation even in conditions of sympatry or incomplete isolation. A robust result is that in an asexual reproduction setting, branching occurs if *σ_α_ < σ_K_*. In the deterministic version of the model, branching will keep occurring indefinitely as a result of the symmetric structure of the fitness landscape (Doebeli 2011). By contrast, stochasticity in the IBM occasionally causes species to go extinct and smoothens the fitness landscape, increasingly slowing branching (see Chap. 3 in Doebeli (2011) for a further description of the process). As a result, the community eventually reaches a stochastic equilibrium. Here, we are primarily interested in determining the speciation rate (that is, the pace of the branching events), as well as the rate of extinction, and how they change with the number of species in the community. This has, to our knowledge, received comparatively little attention. Similarly, while the existence of an equilibrium diversity is known, its value and how it changes with the parameters of the model has not been described, and appears hard to anticipate in a stochastic setting. To help interpreting our results, we approach this quantity empirically by measuring it from the output of the simulations. More precisely, in each simulation, we measured the average diversity at 10 equally distant points in time in the last sixth of total simulation time. We denote the average of this quantity over all replicate simulations in a set by *K*^^^, the “estimated equilibrium diversity”.

### 2.2 Species definition

Although branching in the trait *z* is an emergent feature of the model, species need to be labelled as we do not explicitly model reproductive isolation. Through the course of simulations, we used morphological divergence along the ecological trait axis for translating branching events into speciation events. Specifically, speciation was triggered whenever the two morphologically closest individuals *i* and *j* from the same species were found to diverge in their trait values by more than *θ_z_* (that is, |*z_i_ z_j_| θ_z_*). Upon speciation, we (arbitrarily) assigned the smallest population (cluster) on either side of the morphological gap between *i* and *j* to a new species, or either population at random if both had the same size. In the resulting phylogenies, this had no influence on the length of branches and thus did not affect any of the downstream results. Note that incipient speciation is a feature of the model, as speciation happens some time after two populations start diverging. Incidentally, speciation may fail, for example if the morphological gap collapses as a result of external competition pressures, or if either population goes extinct. However, when speciation does happen, it is permanent: even if the two species become closer in trait value, they are still assumed to be distinct species. Extinction happens when all living individuals of a species fail to produce offspring for the next generation. Note that in our simulations species identity is only a label used to keep track of phylogenetic relationships between individuals and branches, that is, two individuals *i* and *j* respectively belonging to species A and B have strictly the same fitness if *z_i_* = *z_j_*. Importantly, both speciation and extinction are driven by (scramble) competition between individuals: speciation results from branching driven by changes in fitness optima as a result of accumulating local competition intensity, while extinction happens as a result of species being pushed in low-fitness areas of the adaptive landscape.

### 2.3 Simulation procedure

In order to study whether diversity-dependent diversification and the signal it leaves in the phylogenetic tree changes with community size, we ran the LVIBM for an array of values of *σ_K_* (controlling the width of resource distribution and therefore community size) and *σ_α_* (modulating competition intensity, and therefore the number of individuals that can coexist in a community given *σ_K_* (Pontarp et al. 2012)). Specifically, the parameter values we considered were *σ_K_ ∈ {*1, 2, 3, 4, 5*}* and *σ_α_ ∈ {*0.1, 0.2*, …,* 1*}*, resulting in a set of 50 parameter combinations. We set all other parameters to the following values: *r* = 1, *z_opt_* = 0, *K_opt_* = 1000, following Pontarp et al. (2012), and *σ_µ_* = 0.001, *σ_z_* = 0.1. For each parameter setting, we ran 100 replicate simulations (thus 5000 simulations in total). Starting communities contained 10 individuals of the same species, each with trait value *z* = *z_opt_*.

We ran each simulation long enough to allow the community approach equilibrium diversity (here an emergent feature of the model), based on visual examination of the lineage-through-time (LTT) plots of preliminary simulations. In each simulation, we sampled 5% of all individuals in the community at random in every generation where one or more speciation or extinction event happened. For every individual, we saved its generation, its species and the ancestral species it originated from to build a phylogenetic tree of the community.

### 2.4 Estimation of diversity-dependent diversification

We used the phylogenies obtained from the LVIBM to study what form of diversity dependent diversification emerges from competition among individuals. To be consistent with diversity dependent birth-death models (hereafter, DDBD models), we made the assumption that both the rate of speciation (*λ*(*N*)) and the rate of extinction (*µ*(*N*)) are functions of the number of species N. We followed the assumptions made in DDBD models that the rates were not affected by any other factor, such as (generation) time, the affected species’ population size, or the distribution of trait values within it, and that all species in the community were equally likely to be subject to speciation or extinction. For rate reconstruction, we only retained combinations of *σ_α_* and *σ_K_* that produced communities larger than or equal to the arbitrary threshold of 8 species (*K*^^^ 8), trees below this size probably lack the statistical power to yield reasonable estimates of the rates of speciation and extinction. To estimate the rates, we recorded the waiting times to speciation and extinction events and pooled them by the number of species in the community before the event, across all 100 replicate phylogenies for a given setting. We used two methods to estimate the rates from waiting times, both based on maximum likelihood. First, by estimating the rates at each value of *N* separately, maximizing the likelihood of the mean waiting times for each value of *N* . Second, by a standard model selection approach: we assumed a relationship between the rates and diversity, and then estimated the coefficients of this relationship by maximizing the likelihood of the waiting times for all N simultaneously, and finally compared the performance of various relationships specified by the models. We then compared the rates estimated from both approaches.

### 2.5 Estimation for each value of N separately

If we assume that the waiting times for the next event are exponentially distributed we can write down the likelihood for observing, in the simulations, a set of *M* waiting times for each diversity value *N*,

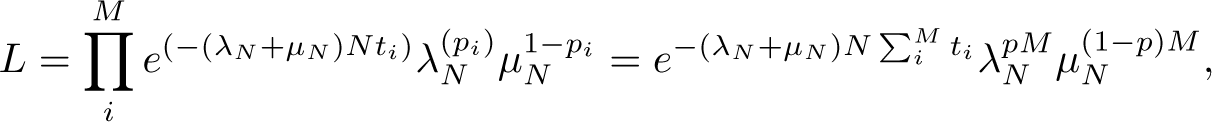

where *p_i_* equals 1 if event *i* is a speciation event and 0 if it is an extinction event, *p* is the proportion of *M* waiting times that lead to a speciation event, *t_i_* is the waiting time until the *i*th event, *λ_N_* is the speciation rate at diversity *N* and *µ_N_* is the extinction rate at diversity *N* . By maximizing the logarithm of this likelihood with respect to both *λ_N_* and *µ_N_*, we can get maximum likelihood estimators for these parameters:

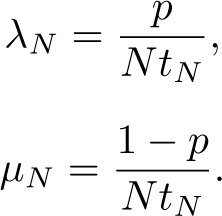

This approach provides estimates of the rates of speciation and extinction corresponding to each value of *N*, and allows us to characterize the main features of the mode of diversity-dependence produced by the LVIBM. However, this approach requires knowledge of the number of species alive in the community through time, and thus is only available for complete trees. We treat the rates estimated from this direct approach as close to the “true” diversity-dependent rates, and use them as a baseline to compare with the rates estimated with the approach below.

### 2.6 Estimation of the coefficients of relationship between the rates and N using birth-death models

To assess what type of diversity-dependent functions were most consistent with the patterns produced by the LVIBM, we maximized the likelihood of the waiting times for all *N* simultaneously for various functions. We considered the linear and power (also known as “exponential” diversity-dependence) functions frequently used in the literature (e.g. Rabosky and Lovette (2008); Etienne et al. (2012)). To investigate the form of diversity-dependence produced by an actual exponential form of diversity-dependence, we also considered an exponential function of the number of species. Diversity-dependence of this form is somewhat intermediate between the linear and power forms, showing asymptotic rates but slower changes in both rates with diversity than in the power function. We used these functions to formulate diversity-dependent speciation and extinction functions (for example, exponential diversity-dependence on speciation, linear diversity-dependence on extinction). In addition, we also considered constant-rate (i.e, diversity-independent) extinction, but not constant-rate speciation, as it is clear from preliminary simulations and previous work using similar IBMs (Doebeli 2011; Pontarp et al. 2012) that speciation slows down as the community grows. Speciation and extinction functions were then paired

together to constitute a set of candidate diversity-dependent models covering all the 12 possible functions (3 speciation functions times 4 extinction functions). We parameterized all diversity-dependent functions to share the same set of four parameters: *λ*_0_, *µ*_0_, *K* and *ϕ*. *λ*_0_ and *µ*_0_ are, respectively the initial speciation and extinction rates when *N* = 0 (or when *N* = 1 in the case of the power functions). *K* is the equilibrium diversity (i.e., the unique value of *N* for which *λ_N_* = *µ_N_*). Finally, *ϕ* controls the value of both rates at equilibrium as a weighted mean of the initial values of both rates,

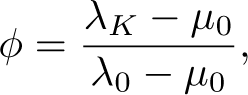

such that

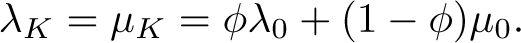

Note that *ϕ* = 0 implies constant-rate extinction, while *ϕ* = 1 implies constantrate speciation. Thus, *ϕ* measures the relative contribution of speciation and extinction to diversity dependence in the model. We bounded 0 *ϕ* 1, and thus assumed that diversity-dependence on speciation is always negative (speciation declines as the community grows), or absent, and that diversity-dependence on extinction is always positive (extinction increases as the community grows), or absent. Below, we refer to this equilibrium rate as *λ_K_*, although this is, by definition, identical to *µ_K_*.

**Table 1:** Speciation and extinction functions used in the birth-death models fitted to the phylogenetic trees produced by the LVIBM.

| Function | Speciation rate | Extinction rate |
| --- | --- | --- |
| Constant | - | $\mu(N) = \mu_0$ |
| Linear | $\lambda(N) = \lambda_0 - (\lambda_0 - \lambda_K) \frac{N}{K}$ | $\mu(N) = \mu_0 + (\lambda_K - \mu_0) \frac{N}{K}$ |
| Power | $\lambda(N) = \lambda_0 N^{-\frac{\log(\frac{\lambda_0}{\lambda_K})}{\log(K)}}$ | $\mu(N) = \mu_0 N^{\frac{\log(\frac{\lambda_K}{\mu_0})}{\log(K)}}$ |
| Exponential | $\lambda(N) = \lambda_0 e^{-\log(\frac{\lambda_0}{\lambda_K}) \frac{N}{K}}$ | $\mu(N) = \mu_0 e^{\log(\frac{\lambda_K}{\mu_0}) \frac{N}{K}}$ |

In all models with constant-rate extinction, we fixed *ϕ* = 0 in the speciation function.

### 2.7 Complete phylogenies

We fitted each of the 12 models to the entire set of 100 replicate complete trees for a given parameter setting of the LVIBM, that is, the entire set of waiting times, for all *N* together, by likelihood maximization.

We used the optimizer function implemented in R package DDD (Etienne et al. 2012), with 1,000 sets of random initial parameter values to minimize the chances of finding only a local likelihood optimum. To restrict the initial values to a range of sensible values, we sampled them in a preset distribution for each parameter. We defined an auxiliary parameter 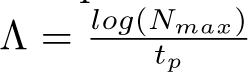 which is an *p* approximation of the average growth rate of the phylogeny assuming constant-rate diversification, with *N_max_* denoting the maximum number of species in the simulation and *t_p_* the simulation time, in generations. We then sampled initial values for *λ*_0_ in uniform distribution *U* (0.5Λ, 2Λ), so that *λ*_0_ was roughly consistent with the size of the tree and the time frame of the simulation. *µ*_0_ was sampled from *U* (0, 0.75*λ*_0_), to avoid a high probability of the tree going extinct early (which can be expected if *λ*_0_ *≈ µ*_0_). Values of *K < N_max_* would make the tree implausible, so we sampled *K* in *N_max_*(1 + Γ(0.5, 0.3)), where Γ is the Gamma distribution, yielding initial values that were most often equal or slightly larger than *N_max_*, with occasionally larger to much larger initial values. Finally, *ϕ* was sampled in *U* (0, 1). For each model, we then selected the parameter values with the largest maximum likelihood out of the 1,000 optimizations, and computed AIC scores and the corresponding AIC weights.

### 2.8 Reconstructed phylogenies

While the model selection approach outlined above gives insight into the type of diversity dependence produced by competition, its results are not directly comparable to an experimental situation where one would seek to estimate diversity-dependence from a single reconstructed phylogeny, where there is no information from extinct lineages. Therefore, we repeated the model selection procedure on single, reconstructed phylogenies in order to estimate how much information is lost as a result of competition-driven extinction. The likelihood for complete trees above cannot be used for reconstructed phylogenies because past diversity in the clade is not directly accessible from the tree. Instead, we used the likelihood and optimization procedure introduced in Etienne et al. (2012) and implemented in the R package DDD. We expanded DDD to include the diversity-dependent functions introduced above that were not already available.

## 3 Results

### 3.1 Diversity-dependence emerges from individual-level competition

Consistent with expectations from adaptive dynamics theory, diversification (i.e., evolutionary branching) occurred in all communities simulated with the LVIBM if *σ_α_ < σ_K_* (that is, all but one setting). Equilibrium community size ranged from a median of *K*^^^ = 2 species to a median of *K*^^^ = 153 (Fig. 2). *K*^^^ is strongly linked to the model parameters: wider resource distributions (larger values of *σ_K_*), and smaller competition kernels (smaller values of *σ_α_*) results in larger communities, with very little variability across replicates (Fig. 2). We note, however, that not all communities completely reached equilibrium diversity by the end of simulations. Equilibrium diversity, as measured by *K*^^^, appears to be entirely predictable from the values of parameters *σ_K_* and *σ_α_* (Fig. 2).

**Figure 1:**
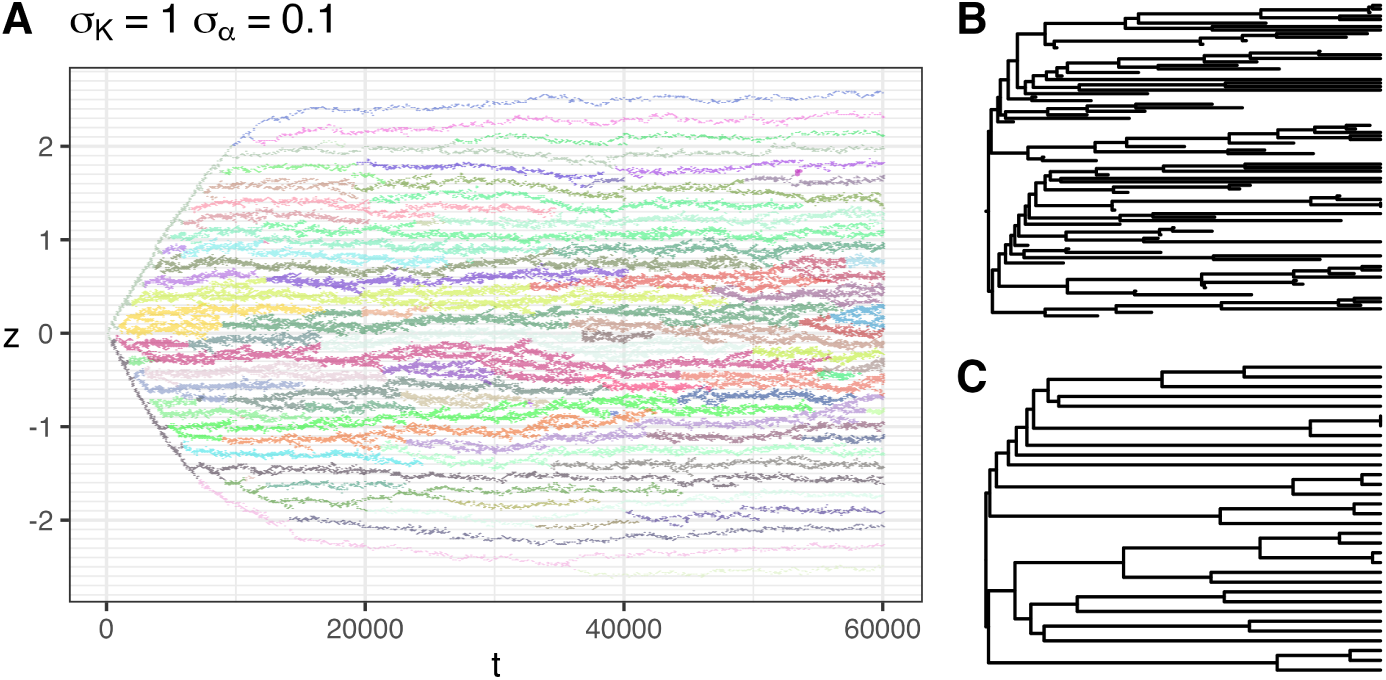
Example output of the individual-based model for parameters *σ_K_*= 1, *σ_α_* = 0.1, displaying diversification. (A) Sample of the distribution of phenotypic clusters (species) in trait space over 60,000 generations. 5% percent of all individuals in the community were sampled every 200 generations, and coloured by their species identity. (B) Complete and (C) reconstructed phylogenies built from the community displayed in (A).

**Figure 2:**
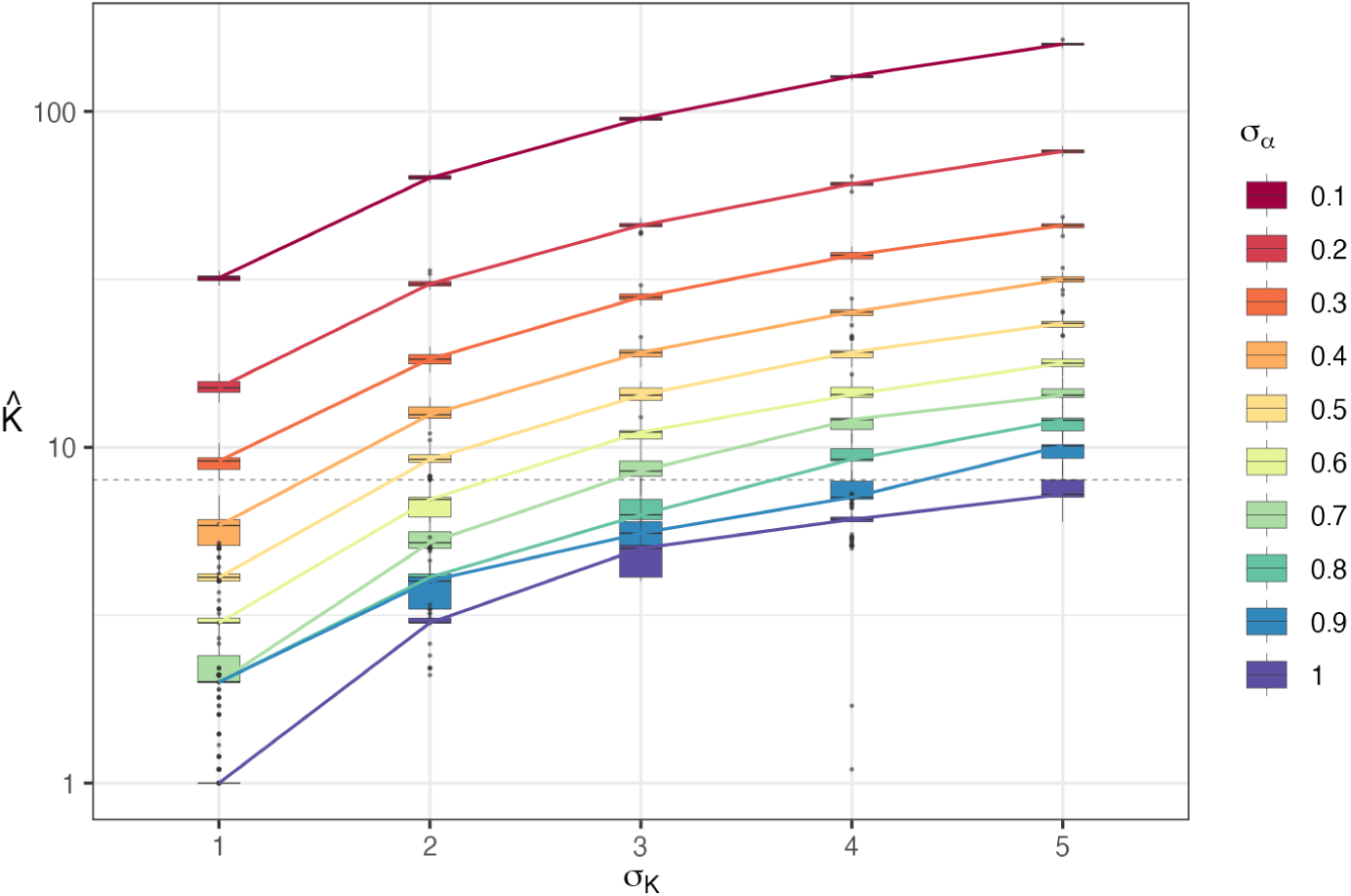
Estimated equilibrium diversity (*K*^^^) at the end of simulations from communities simulated with a range of resource abundance widths (*σ_K_*) and competition kernel widths (*σ_α_*), shown on a log10-scale. Boxplots represent the distribution of *K*^^^ for each of the 100 replicates from the same parameter set, with lines representing the medians for each parameter set. Parameter sets with a median equilibrium diversity under *K*^^^ = 8 species (dashed line) were discarded from subsequent analysis.

We found *K*^^^ is proportional to the ratio between *σ_K_*, which controls the*_√_*t<u>ot</u>al amount of phenotype space (through the total area under *K*(*z*), *σ_K_K_opt_* 2*π*), and *σ_α_*, which controls the width of low-fitness gaps between branches (and thus, how many species can coexist under *K*(*z*) while maintaining non-negative growth). We also note that *σ_α_* also affected *K*^^^ negatively independently beyond this ratio, although we could not establish the exact relationship linking *K*^^^ with the two parameters. This is consistent with the results of Doebeli and Ispolatov (2017) in the context of a multidimensional *K*(*z*); they reported the equilibrium diversity to scale exponentially with the dimensionality of *K*(*z*) and to decrease with increasing *σ_α_*. We found that fair predictions of *K*^^^ can be obtained in our results with the empirical relation,

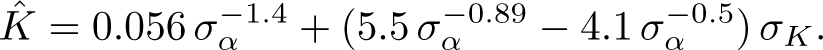

This relation is not entirely satisfactory however, as it does not capture important predictions from adaptive dynamics: *K*^^^

should equal 1 if *σ_α_* = *σ_K_*, and the number of species should increase towards infinity as *σ_α_* approaches zero, independently of the value of *σ_K_*. Diversification is definitely diversity-dependent in our simulations. Species-(Fig. 3) and lineage-through-time (Fig. 4) plots of the phylogenies built from simulated communities have a shape consistent with diversity-dependent diversification, with initially fast branching followed by a slowdown of diversification, and the eventual saturation of the community around *K*^^^ (see Etienne et al. (2012); Pannetier et al. (2021) for comparison with phylogenies simulated under a diversity-dependent birth-death model). The average rate of speciation across replicate communities, as measured from the mean waiting time between successive events, decreases with species diversity, while the extinction rate tends to increase (Fig. 5, Fig. 6). The restricted variation in tree size across replicates (Fig. 2) is unusual for a diversity independent process, and instead supports a structuring effect as induced by the feedback between diversity and diversification in diversity-dependent models (Pannetier et al. 2021).

**Figure 3:**
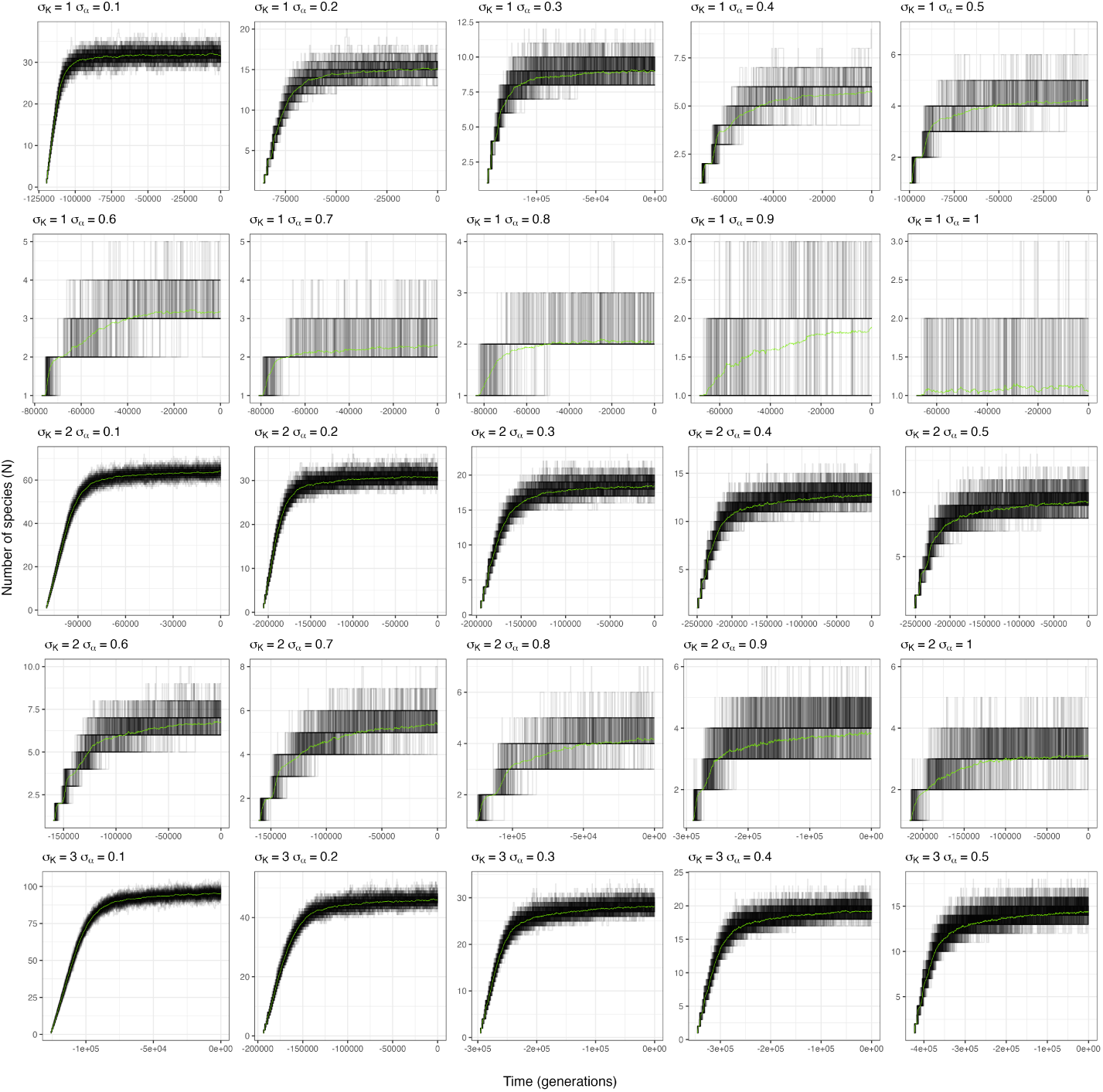
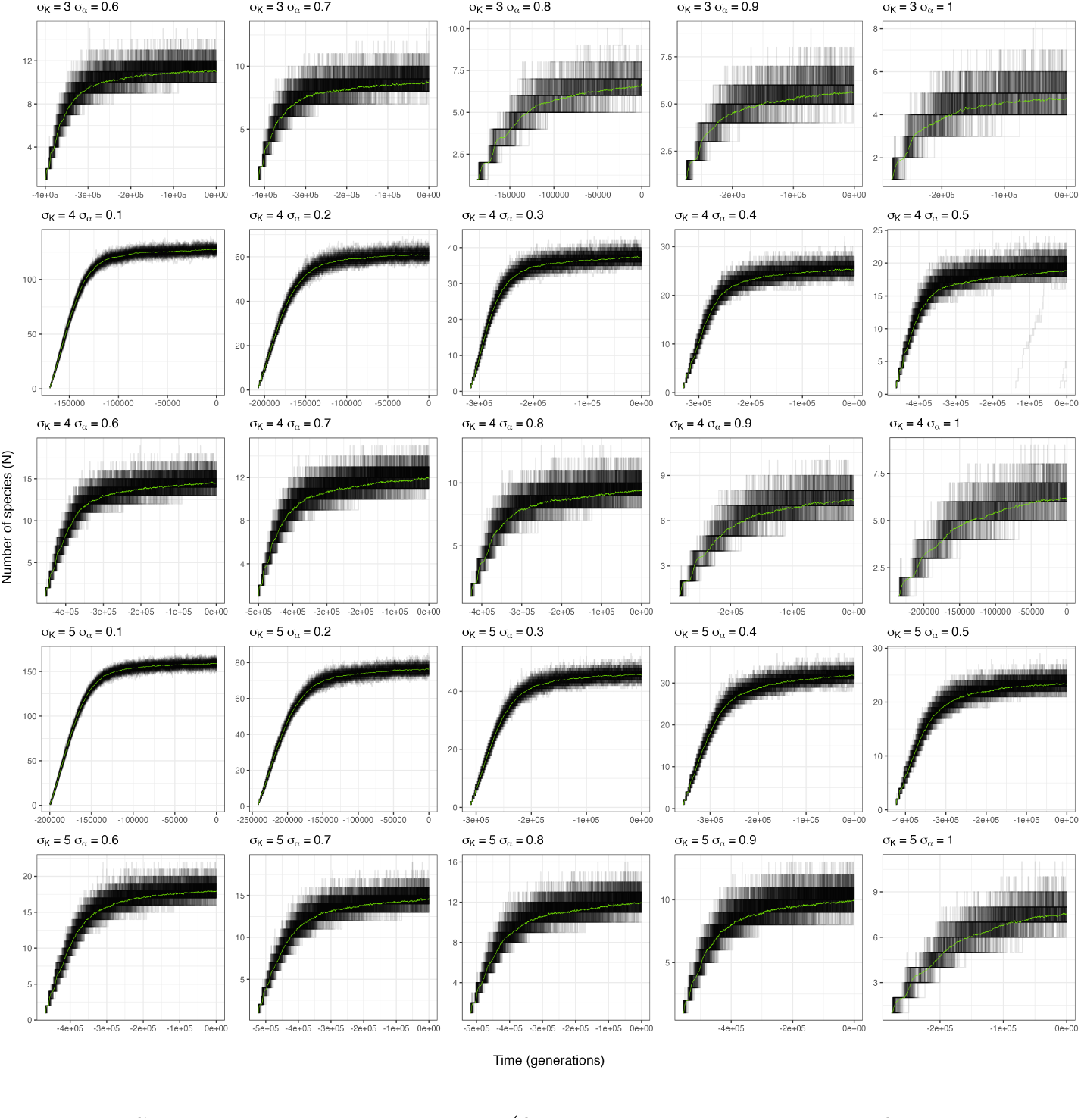
Species-through-time plots (STT, that is the number of species in the clade at a given time in the past) of the phylogenies produced by the LVIBM. Each panel displays the individuals STTs for each of the 100 replicate simulations (black lines), and the average STT (green curve), computed at 1,000 equidistant points in time, for a given parameter setting.

**Figure 4:**
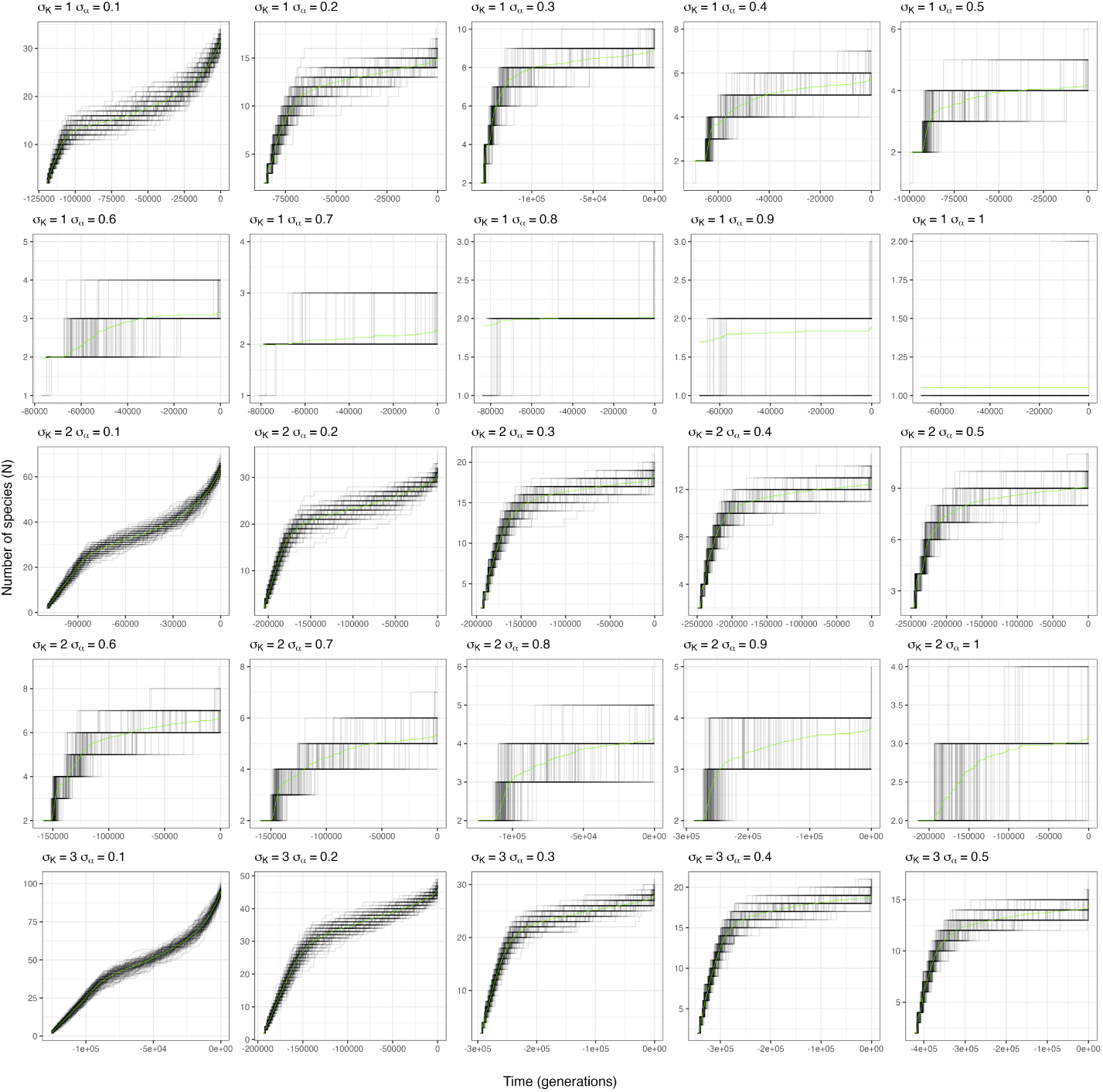
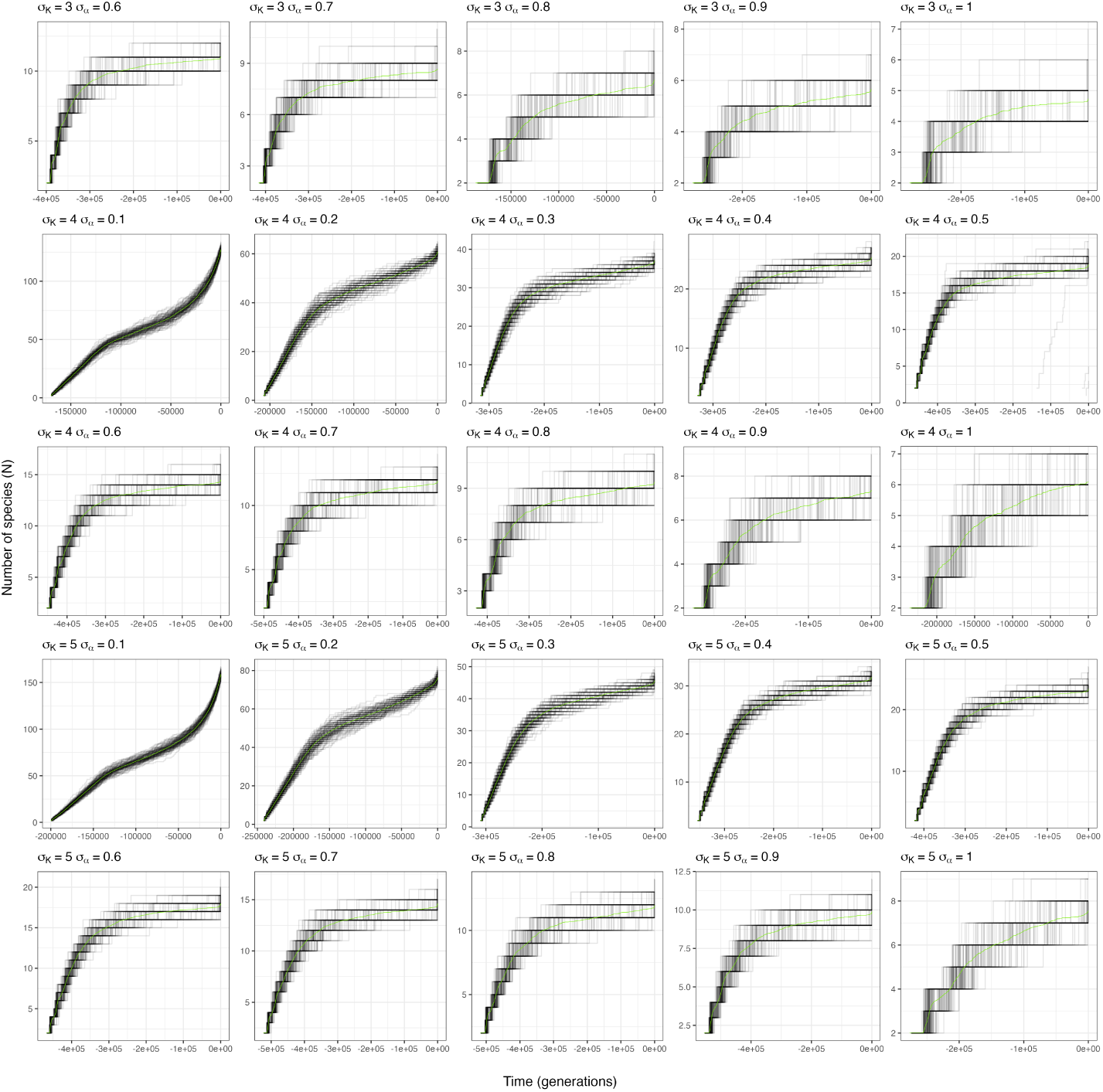
Lineage-through-time plots (LTT, that is the number of lineages at a given time in the past that have extant descendants at the present) of the phylogenies produced by the LVIBM. Each panel displays the individuals LTTs for each of the 100 replicate simulations (black lines), and the average LTT (green curve), computed at 1,000 equidistant points in time, for a given parameter setting.

**Figure 5:**
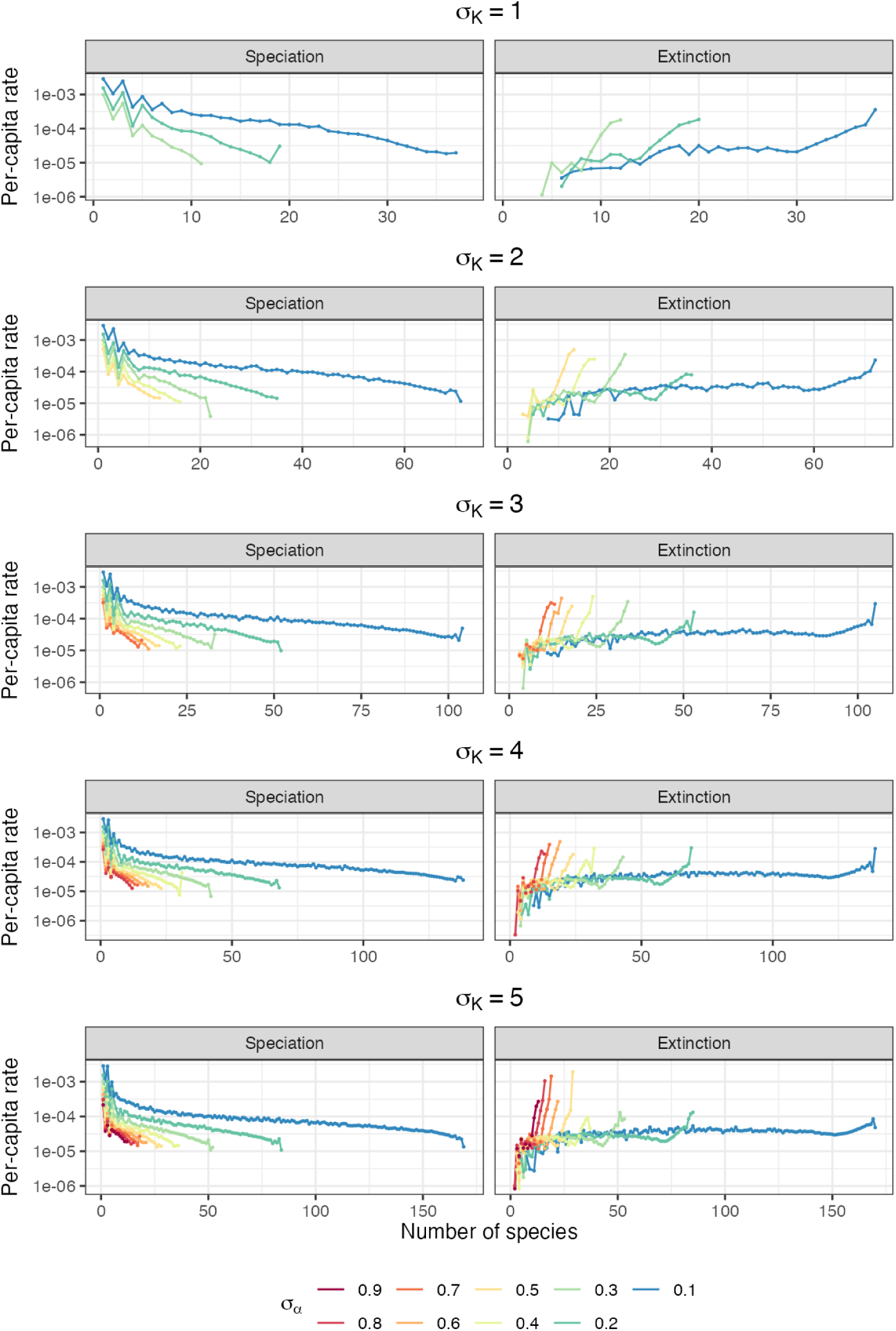
Per-capita rates of speciation and extinction across values estimated independently for each value of *N*, from the inverse of the mean waiting time between events. Note the log10-scale of the y-axis.

**Figure 6:**
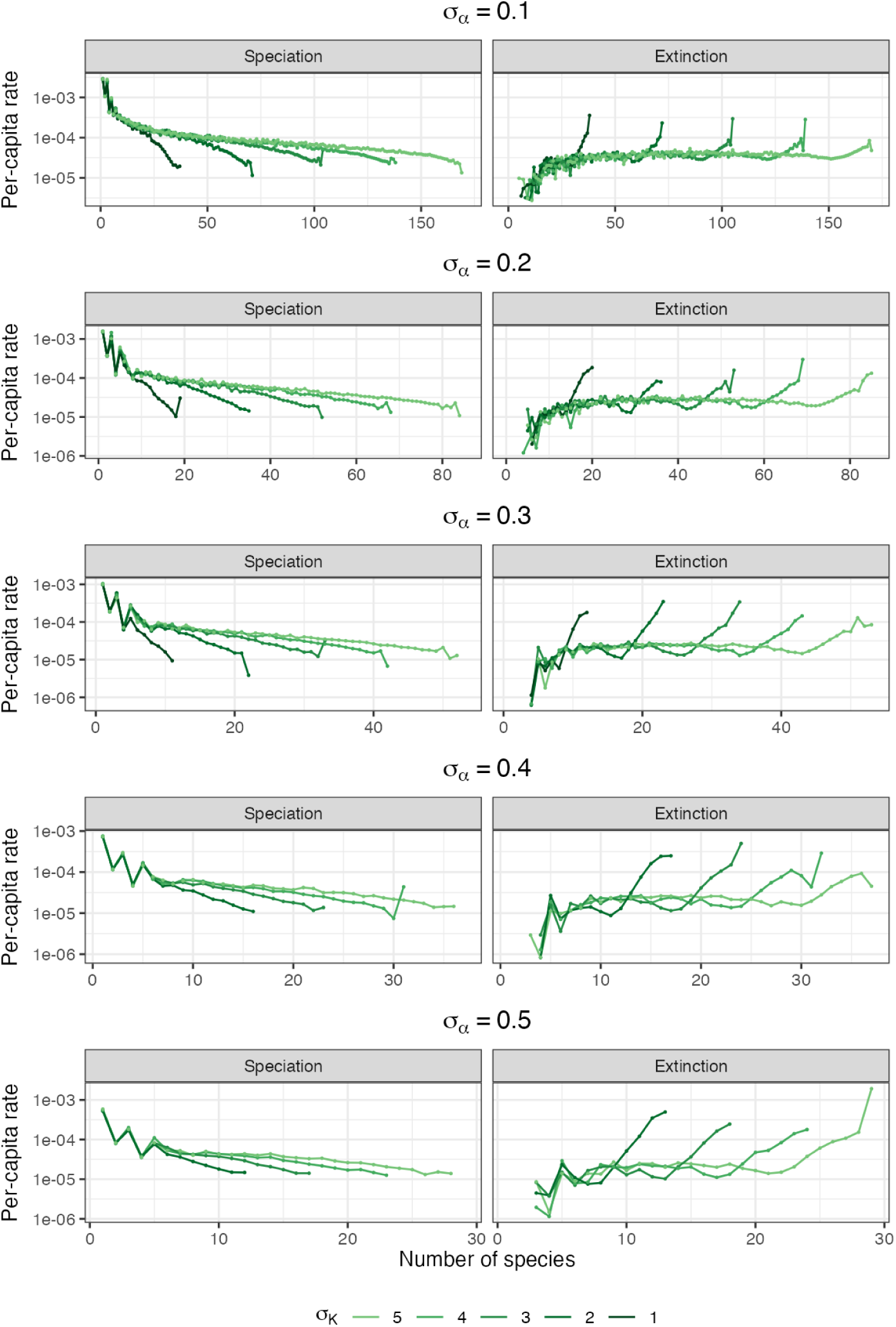
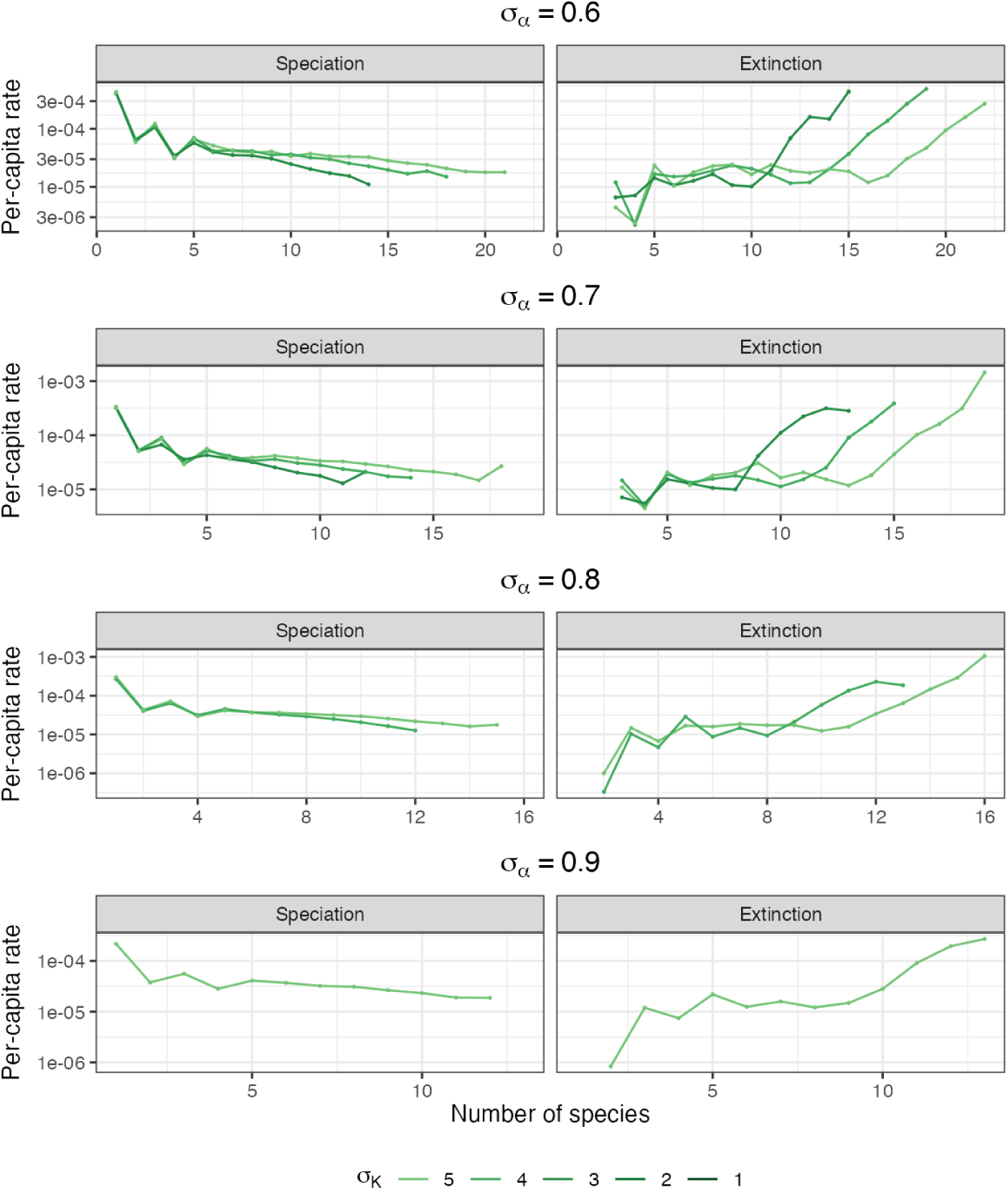
Per-capita rates of speciation and extinction across values estimated independently for each value of *N* . The values are the same as in Fig.5, but are displayed by value of *σ_K_* instead of *σ_α_*.

The relationship between species diversity and the per-capita rates of speciation and extinction is more complex than a simple increase or decrease (Fig. 5, Fig. 6). The decline of speciation progresses in two phases. Speciation is initially very high and decreases quickly with the number of species. During this first phase, the speciation rate oscillates as it declines (Fig. 5, Fig. 6). This is a result of the synchronous nature of this initial phase of the diversification process: branching tends to occur simultaneously on both crown lineages (Fig. 1), causing rapid speciation from e.g. 2 to 4 species. Such simultaneous branching of crown lineages is typical of the deterministic version of the model (Doebeli 2011). Oscillations dampen after the few first branching events as stochasticity causes speciation events to decouple (Fig. 1). Remarkably, the average rate of speciation in this first phase is identical across all values of *σ_K_* (but not *σ_α_*, Fig. 5, Fig. 6). Then, the decline of speciation with diversity itself slows down, and the speciation rate enters a second phase where it keeps declining at a constant pace with the number of species. The slope of the decline in that second phase depends on the value of *σ_K_* and *σ_α_*, with low values of *σ_K_* and high values of *σ_α_* being associated with a steeper slope (Fig. 5, Fig. 6). Low *σ_K_* limits the width of exploitable trait space, while high *σ_α_* pushes species further apart from one another in trait space. Both thus contribute to limiting the number of species that can coexist at equilibrium, and increasing the intensity of diversity-dependent effects on speciation. In addition, *σ_α_* also affects the absolute value of the speciation rate: higher competition intensities results in an overall lower speciation rate (Fig. 5, Fig. 6). Extinction is initially absent, and only starts occurring after a few species have branched (Fig. 5, Fig. 6). On its onset, the rate of extinction initially increases very fast, but this fast increase also slows down quickly as more species accumulate in the community. Upon diversity reaching a certain value, the extinction rate starts increasing quickly again, exceeding the speciation rate (and thus, setting equilibrium diversity), and keeps increasing steeply beyond this point.

Between these two phases, we observe an intermediate phase of diversity dependence where the extinction rate is about constant, or follows a hump, increasing at a slower pace than previously, then decreasing with the number of species shortly before the start of the second phase of acceleration of extinction. Interestingly, this decrease in the extinction rate appears to match the pace of the decrease in the speciation rate, such that the net rate of diversification (speciation - extinction) in this phase is about constant. This intermediate phase is hard to observe and may not be present in the smallest communities, where extinction instead seems to increase without interruption, but in all communities with more than about 20 species, the occurrence of extinction at a constantrate is clear. This intermediate phase is only visible in settings producing large communities. For smaller communities, only a continuous, steep increase of the extinction rate is visible, as equilibrium diversity is reached quickly after the initial onset of extinction. Neither *σ_K_* nor *σ_α_* appear to have an effect on the absolute rate of extinction, or the extent of its increase, although this is not clear due to the relatively small number of extinction events at low levels of diversity (Fig. 5, Fig. 6). Both parameters appear to have an effect on how early extinction started to increase, and the duration of the different phases, through their effect on equilibrium diversity.

Thus, despite important quantitative differences in the diversity-dependent rates across parameter settings, the form of diversity-dependence emerging from the LVIBM can be described as a single process, with parameters *σ_K_* and *σ_α_* controlling (through equilibrium diversity) the diversity at which the process transits from one form to the next. In summary, *σ_K_* and *σ_α_* modulate the speciation and extinction rates through their effect on setting the equilibrium diversity of the community: higher values of *σ_K_* increase it, while higher values of *σ_α_* decrease it. This in turn influences the slope of the speciation rate, and the diversity at which the extinction rate accelerates towards infinity (Fig. 5, Fig. 6). Independently of the equilibrium diversity, smaller values of *σ_α_* also increase the speciation rate overall, while no such effect is visible on the extinction rate.

### 3.2 Extinction mostly concerns exclusion between sister lineages

It has been suggested that diversity-dependence happens through competitive exclusion, a view consistent with Darwin’s own proposed process of diversification, and that thus has been referred to as Darwinian diversity-dependence (Rabosky 2013). In our simulations, we do observe that species are at a much higher risk of extinction after branching: the distribution of the length of branches leading to extinction is strongly skewed to the left (Fig. 7). This concerns both the new daughter lineage appearing with the speciation event, and the mother lineage undergoing speciation. Extinction events concerning branches younger than 500 generations concern over 20% of all extinction events, in all parameter settings (Fig. 7). Competition intensity has an important effect on the proportion of these early extinction events, which made up around 20% of all extinction events as stated above for *σ_α_* = 0.1, and up to 56-57% in settings with *σ_α_ >* 0.6. We examined the contribution of these short branches to diversity-dependence in the extinction by observing how the proportion of short-branch extinction events (out of all extinction events) changes with the value of *N* (Fig. 8). Overall, changes in the proportions have a small amplitude, staying about constant for much of the range of values of *N* . A continuous increase of the proportion of short-branch extinctions with increasing diversity is visible for settings with *σ_α_* 0.2. In those cases, short-branch extinction represents a very low proportion of extinction events at low diversity, suggesting that competition is sparse enough for allowing both sister lineages to survive after divergence. For settings with *σ_α_ >* 0.2, the pattern is instead U-shaped: short-branch extinctions represent the majority to totality of all extinctions when *N* is low. Excluding short branches from the data decreases both the speciation and extinction rate significantly (Fig 9). Yet, we observe that diversity-dependence remains largely unaffected: changes in both rates with values of *N* are virtually identical whether short-lived species are excluded from data or not. In summary, while a significant part of all extinction events concerns the extinction of recently diverged species as a result of competitive exclusion by their sister lineage, the proportion these events represent is fairly uniform across diversity bins, and thus does not contribute to diversity-dependence substantially.

**Figure 7:**
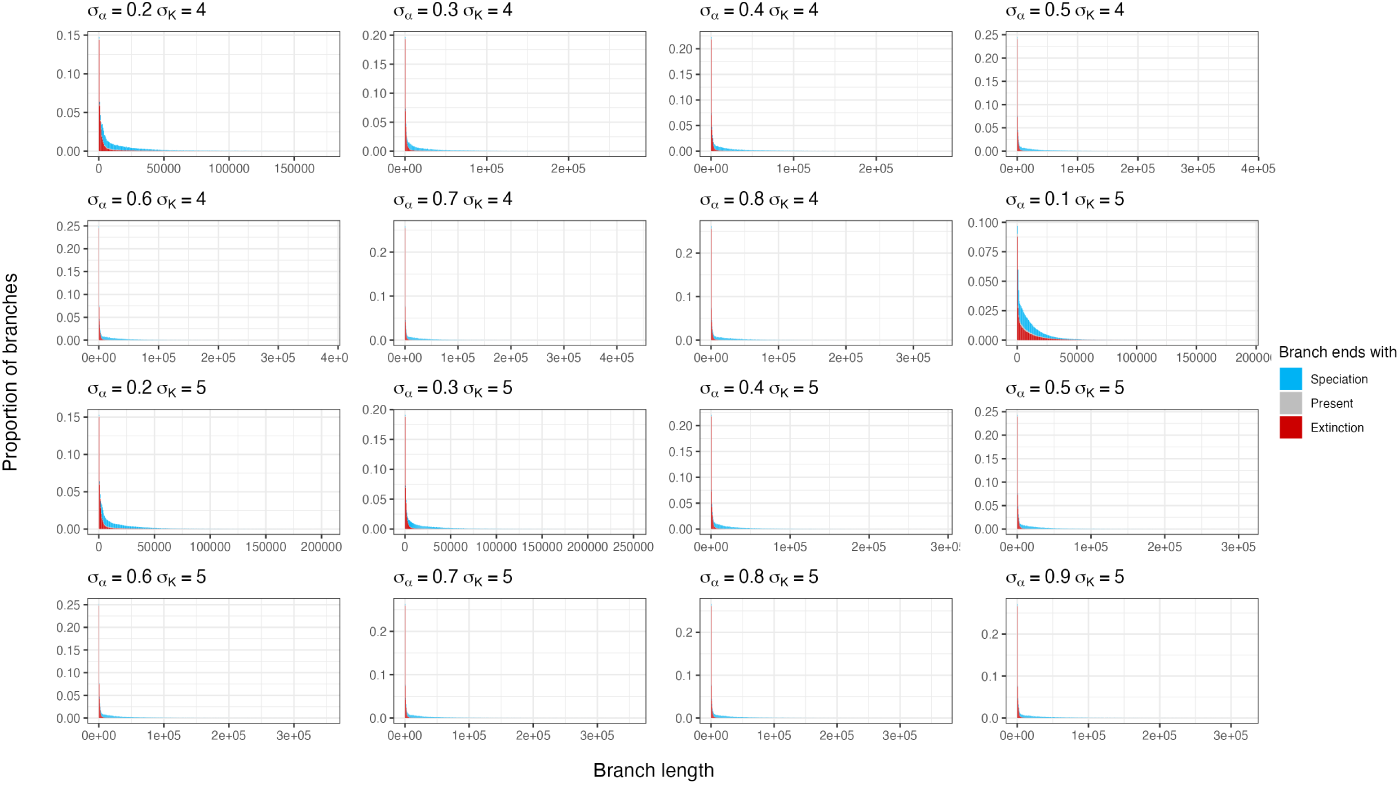
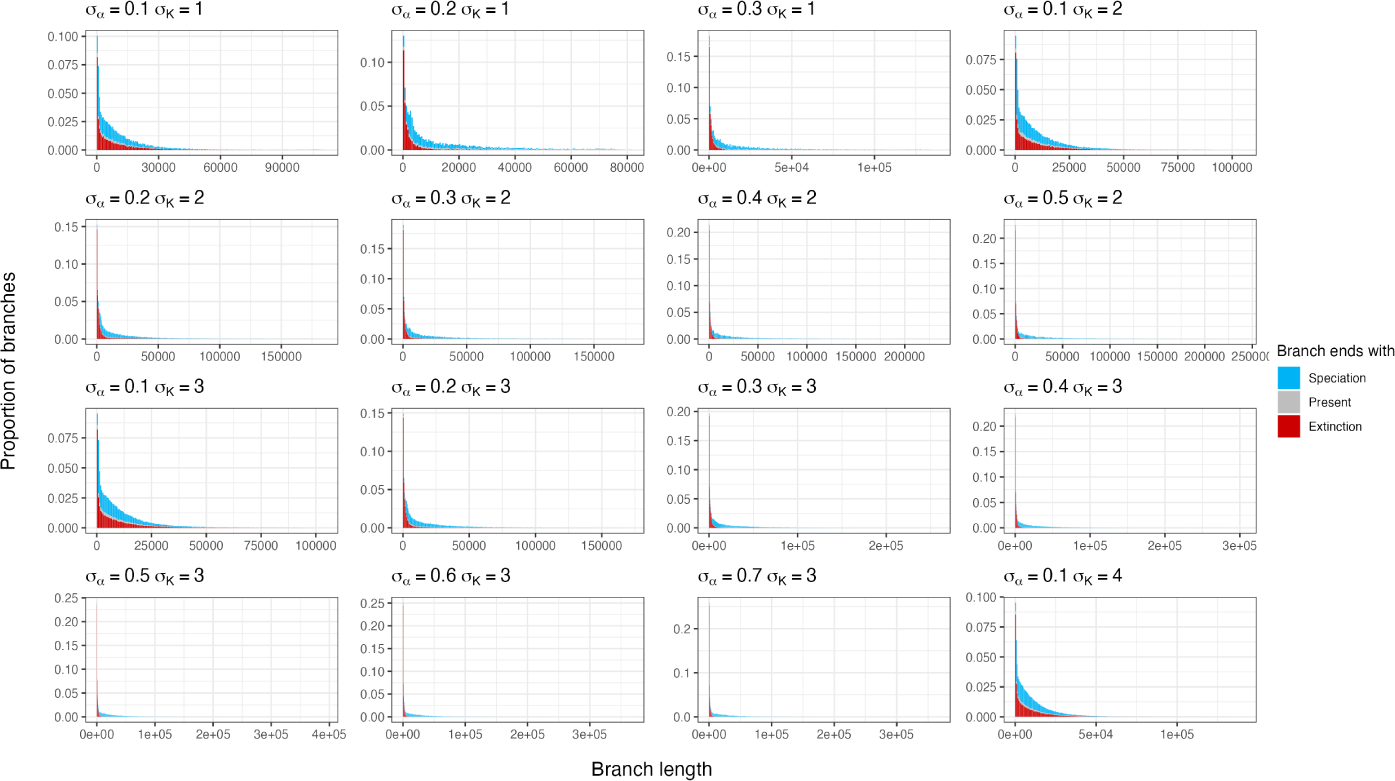
Distribution of the length of branches appearing at a given point in time, coloured by the event terminating the branch.

**Figure 8:**
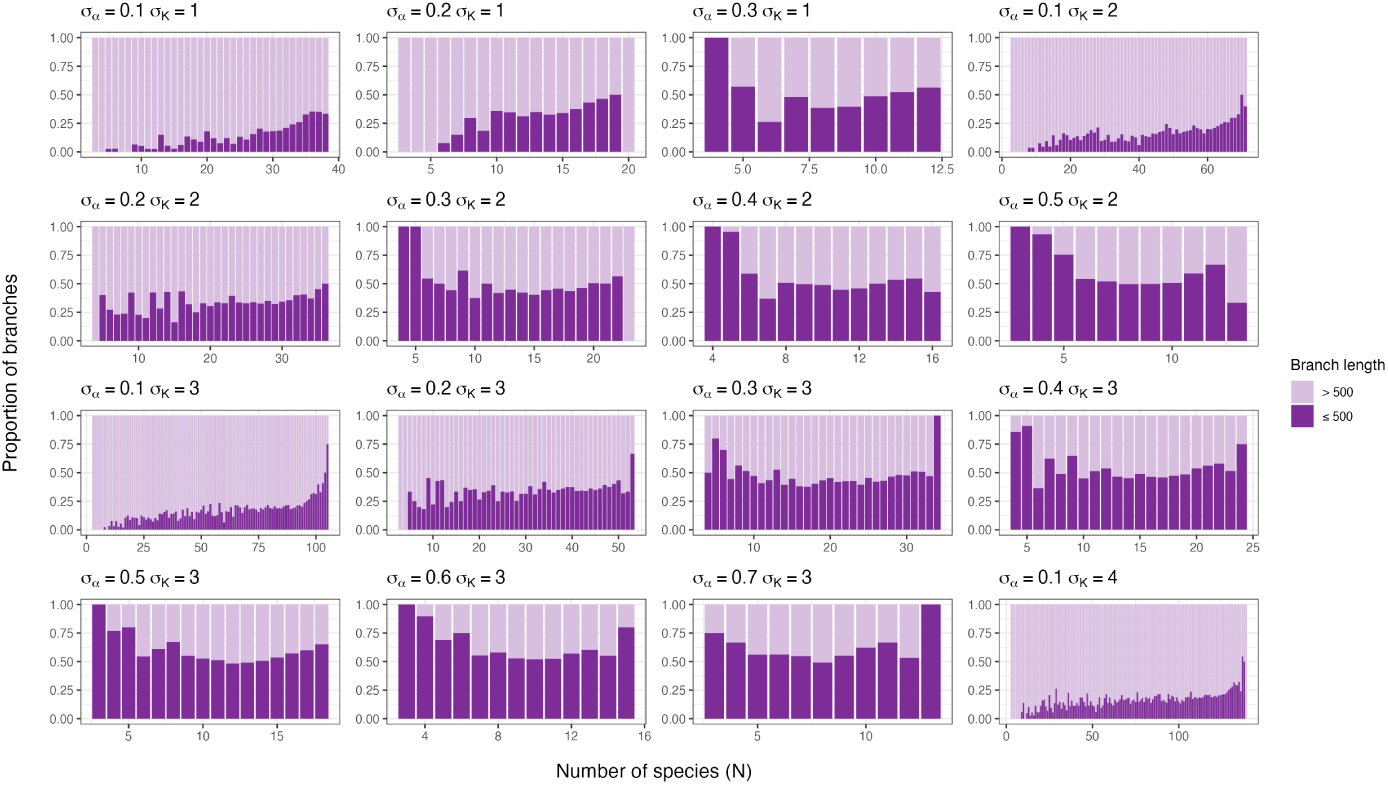
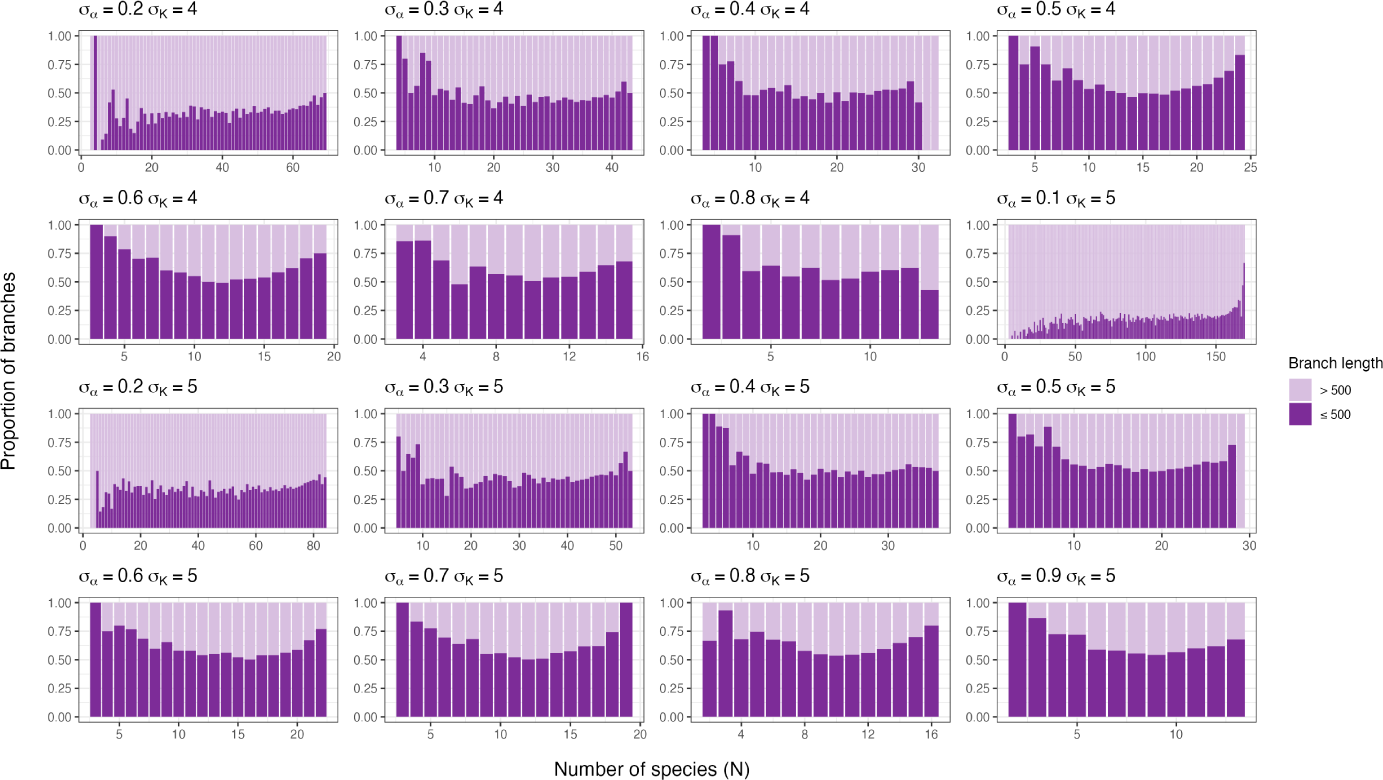
Proportion of short branches (*<* 500 generations) among branches leading to extinction, plotted as a function of *N* .

**Figure 9:**
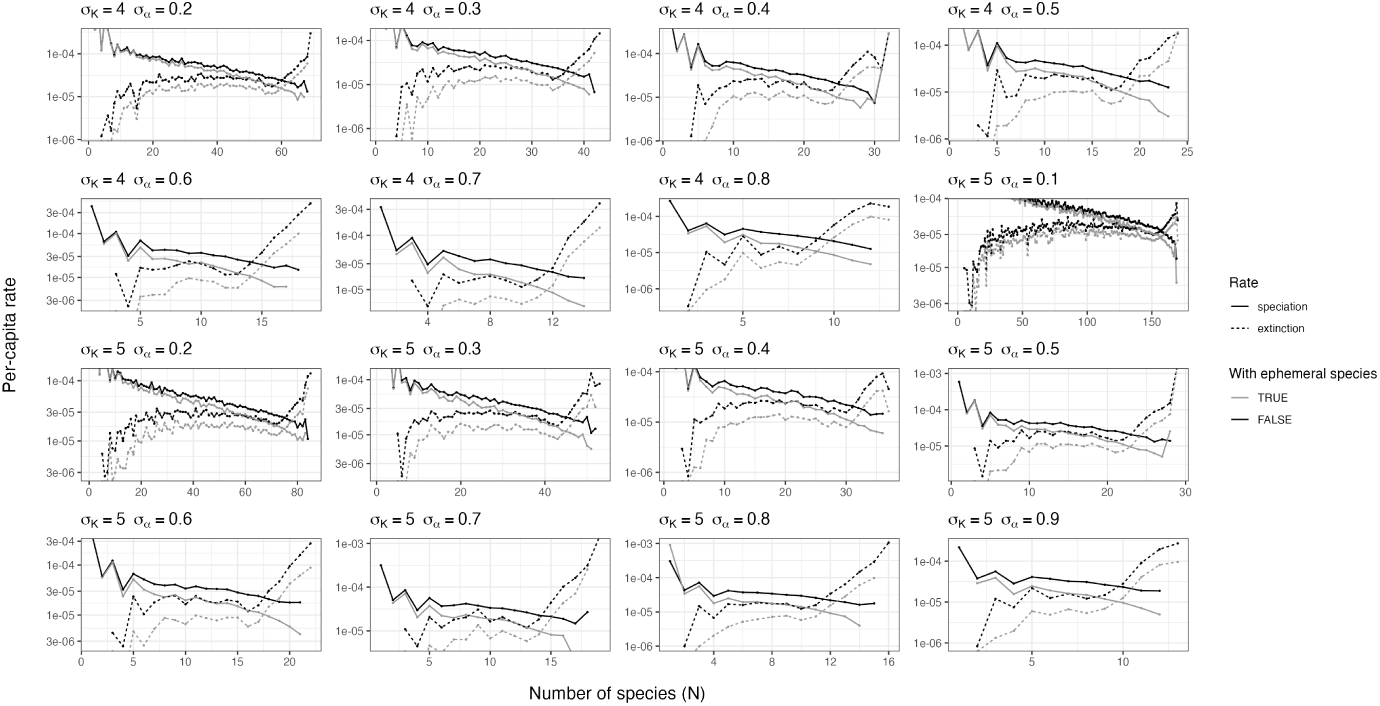
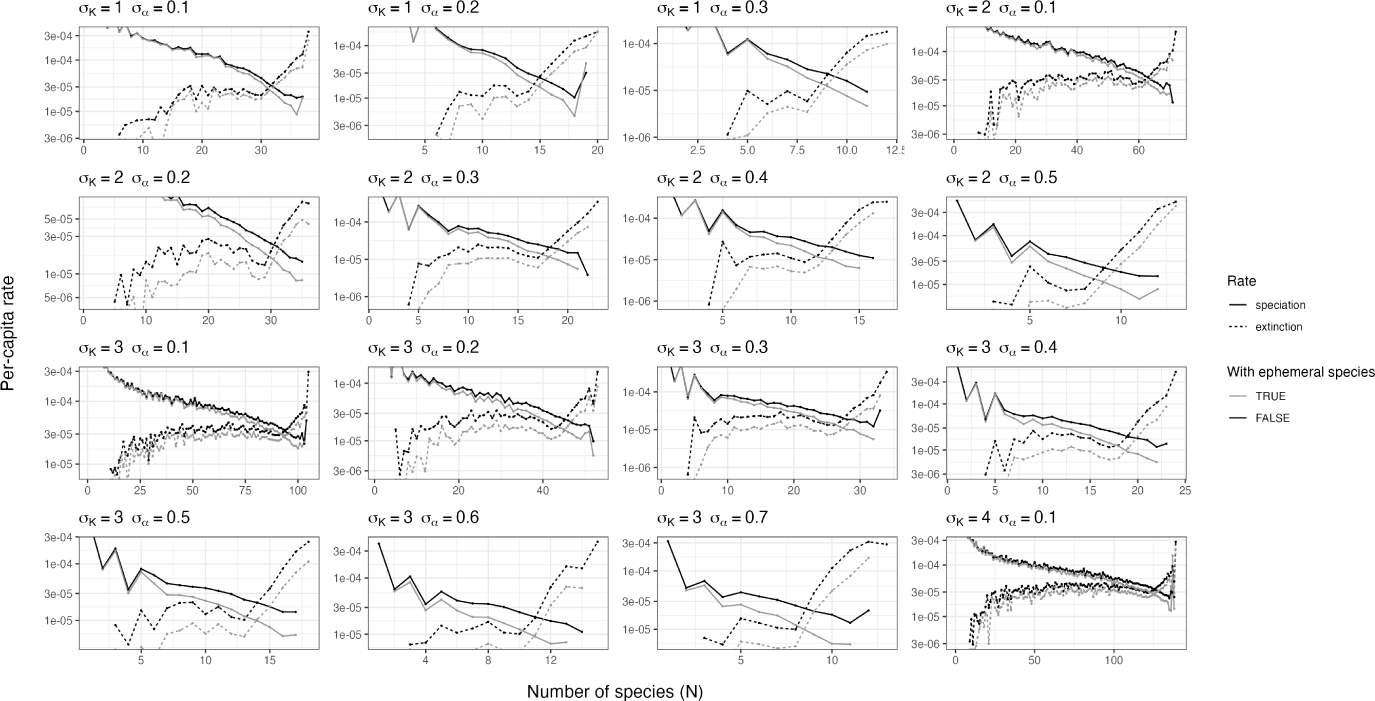
(continued) Per-capita rates of speciation (full lines) and extinction (dashed lines), estimated independently for each value of *N*, including (black lines) or excluding (grey lines) ephemeral species (defined as species that survive for less than 500 generations). Black lines are the same curves as in Fig. 5 and Fig. 6

### 3.3 Model selection on complete trees

In all settings, one single model was decisively supported over all others, although this best model itself varied across parameter settings. The AIC weight for the best model was over 0.95 in all but one setting (*σ_K_* = 4, *σ_α_* = 0.2 where *AIC_w_* = 0.71 for the best model) (Fig. 10). Rather than a much-improved performance in approximating the LVIBM from the best model compared to others, these large scores appear to be a result of the large number of observations in the data (i.e., all waiting times between successive events across 100 replicate simulations, 9.103 to 9.105 observations) and the small variability across replicates (Fig. 2). Reducing the number of observations to solve this issue would make little sense, as *AIC_w_* scores would then reflect sample size rather than the relative performance of each model. Below, we interpret the selected model as providing the best approximation, but rely on inspection of the rates predicted by each function to interpret how well each type of diversity-dependence approximates the rates emerging from the LVIBM.

**Figure 10:**
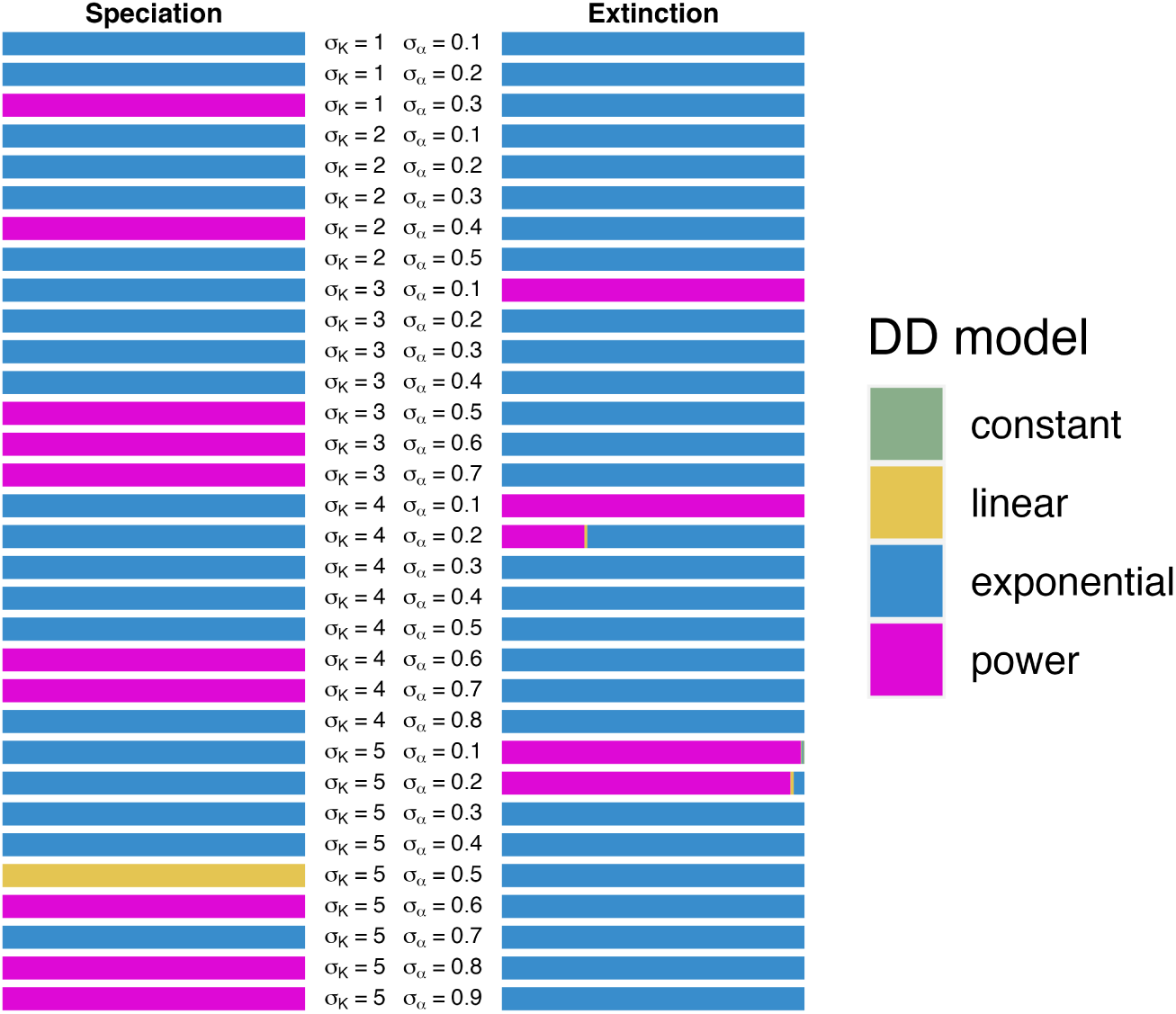
Results of model selection using birth-death models and maximum likelihood, for complete trees. The width of the bars denotes the total AIC weight for each diversity-dependent function, taken as the sum of AIC weights of all birth-death models that included that function.

#### 3.3.1 Exponential- or power diversity-dependence best approximates the form of diversity-dependence emerging from the LVIBM

For all but one combination of *σ_K_* and *σ_α_*, the best fitting model contains either exponential or power diversity-dependence on the speciation rate, and either exponential or power diversity-dependence on the extinction rate (Fig. 10). Linear diversity-dependence in the speciation rate was supported in only one setting (*σ_K_* = 4, *σ_α_* = 0.5), while constant-rate extinction was never supported (Fig. 10). The selection of exponential or power diversity dependence in the speciation rate appears to be driven by the early, explosive phase of speciation that can be observed from the rates estimated separately (black lines in Fig. 11). Explosive speciation was indeed absent from the speciation rate predicted by the linear model (yellow lines in Fig. 11). This pattern was instead best approximated by power diversity-dependence, and to a lesser extent, by exponential diversity-dependence (Fig. 11, pink and blue lines, respectively). This early phase appears to have a strong influence on the differences in likelihood between the models, as power diversity-dependence otherwise tended to consistently underestimate the rate of speciation compared to linear diversity-dependence, particularly in the largest communities. The selection of exponential diversity-dependence seems to be the result of this model providing estimates intermediate between power and linear diversity-dependence. For the smallest communities (*K*^^^ *<* 20 species), estimates of the speciation rate after the initial phase are closer to those of other models, which may explain why power diversity-dependence in speciation tends to be selected over exponential diversity-dependence in these cases (Fig. 10).

**Figure 11:**
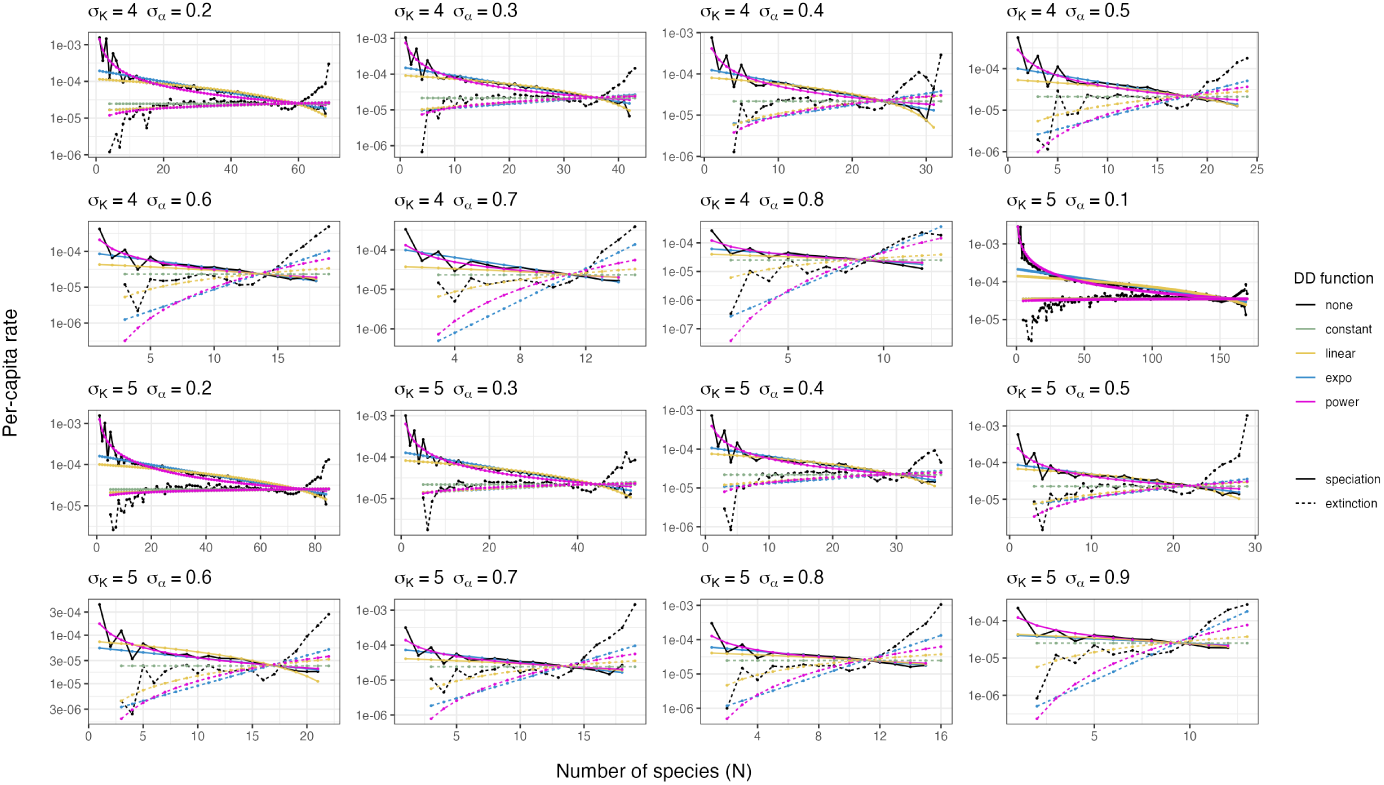
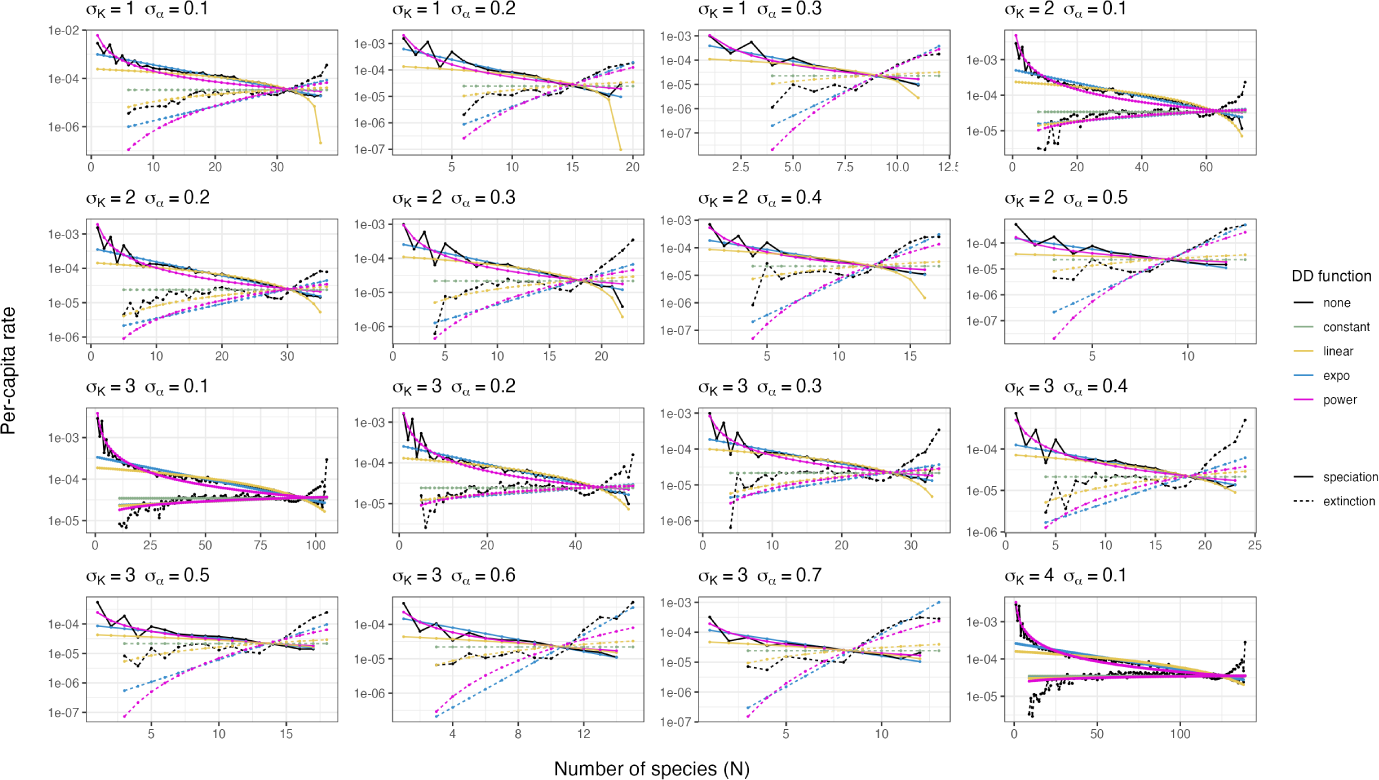
Maximum-likelihood estimates of the speciation and extinction rates obtained from complete trees (coloured lines). Rates estimated independently for each value of *N* (Fig. 6) are plotted as well (black lines) for comparison.

Exponential diversity-dependence in the extinction rate was selected for all parameter settings except the four that produced the largest communities (Fig. 10), where power diversity-dependence was instead preferred. Here again, the selection of these two models over linear diversity-dependence and constant-rate extinction appears to result from the rapid (rather than gradual) onset of extinction. Without this feature, linear diversity-dependence, or constant-rate extinction appears to better approximate the extinction rate observed in the LVIBM (Fig. 11), suggesting that model selection is strongly influenced by these early extinction events. This is particularly visible in the largest communities, where power diversity-dependence is selected over constant-rate extinction despite the extinction rate being close to constant for a large part of the diversification process (see previous section).

The shared features of power and exponential diversity-dependence allow highlighting important aspects of the mode of diversity-dependent diversification that emerges from competition at the level of individuals: the speciation rate decreases, and the extinction rates decrease with the number of species. The decline of speciation and onset of extinction are initially steep, but the rate of change itself declines quickly with the number of species, although it did not reach zero for the range of values of *N* covered in the community data.

#### 3.3.2 Diversity-dependence is mostly mediated by the speciation rate

Estimates of parameter *ϕ* are well below 0.5 for most parameter settings, indicating a more important contribution of the speciation rate than the extinction rate to overall diversity-dependence in diversification, although the value of *ϕ* increased with higher values of *σ_α_* (Fig. 15). That is, for settings with low competition intensity (*σ_α_* = 0.1, 0.2), *ϕ* was estimated as close to zero, such that the extinction rate changed little with the number of species compared to the speciation rate. Intriguingly, in such settings, estimates of *ϕ* for models containing exponential or power diversity-dependence in the speciation rate lie close to zero, suggesting nearly-constant extinction. This is consistent with the rates estimated separately for each *N* : in settings with low *σ_α_*, extinction tends to change very little over intermediate values of *N* (Fig. 6). Yet, the rates associated with these estimates do differ from constant-rate extinction by an initial rapid increase (Fig. 11), and this appears to be sufficient to prompt their selection over constant-rate extinction. In communities with high competition intensity (e.g, *σ_α_* = 0.7, 0.8, 0.9), by contrast, estimates of *ϕ* are higher, and in some instances, close to 0.5 (Fig. 15), indicating that increasing *N* increases the extinction rate about as much as it decreases the speciation rate. Again, this is consistent with the rates estimated separately (Fig. 10 and black lines on Fig. 11), where the fast saturation of niche space resulted in a steep, uninterrupted increase of the extinction rate from its onset. To summarise, the same evolutionary scenario (i.e., the LVIBM) leads to a varying degree of diversity-dependence in the speciation rate versus extinction rate. In general, diversity-dependence affects the speciation rate more than the extinction rate. Yet if equilibrium diversity is low enough to be reached quickly, diversity-dependence affects both rates more evenly.

### 3.4 Model selection on reconstructed trees

#### 3.4.1 Model selection recovers linear diversity-dependence in the speciation rate

We find that the exponential and power diversity-dependence in the speciation rate are not recovered in reconstructed trees. Instead, linear diversity-dependence was decisively supported by average AIC weights across all settings (Fig. 12). Contrary to the case of complete trees, the large support observed cannot be attributed to the large size of the datasets (models were fitted to each tree separately). Summarising support with an alternative method, by counting the number of occurrences across replicates of each model being selected as the best model yields even stronger support for linear diversity-dependence in speciation (Fig. 13). This is in contrast with the very strong support for exponential and power diversity-dependence found in the case of complete trees, and suggests that the strong signal for initially explosive speciation that appears to drive the fit of the model in the complete tree case is lost with extinct lineages in reconstructed trees.

**Figure 12:**
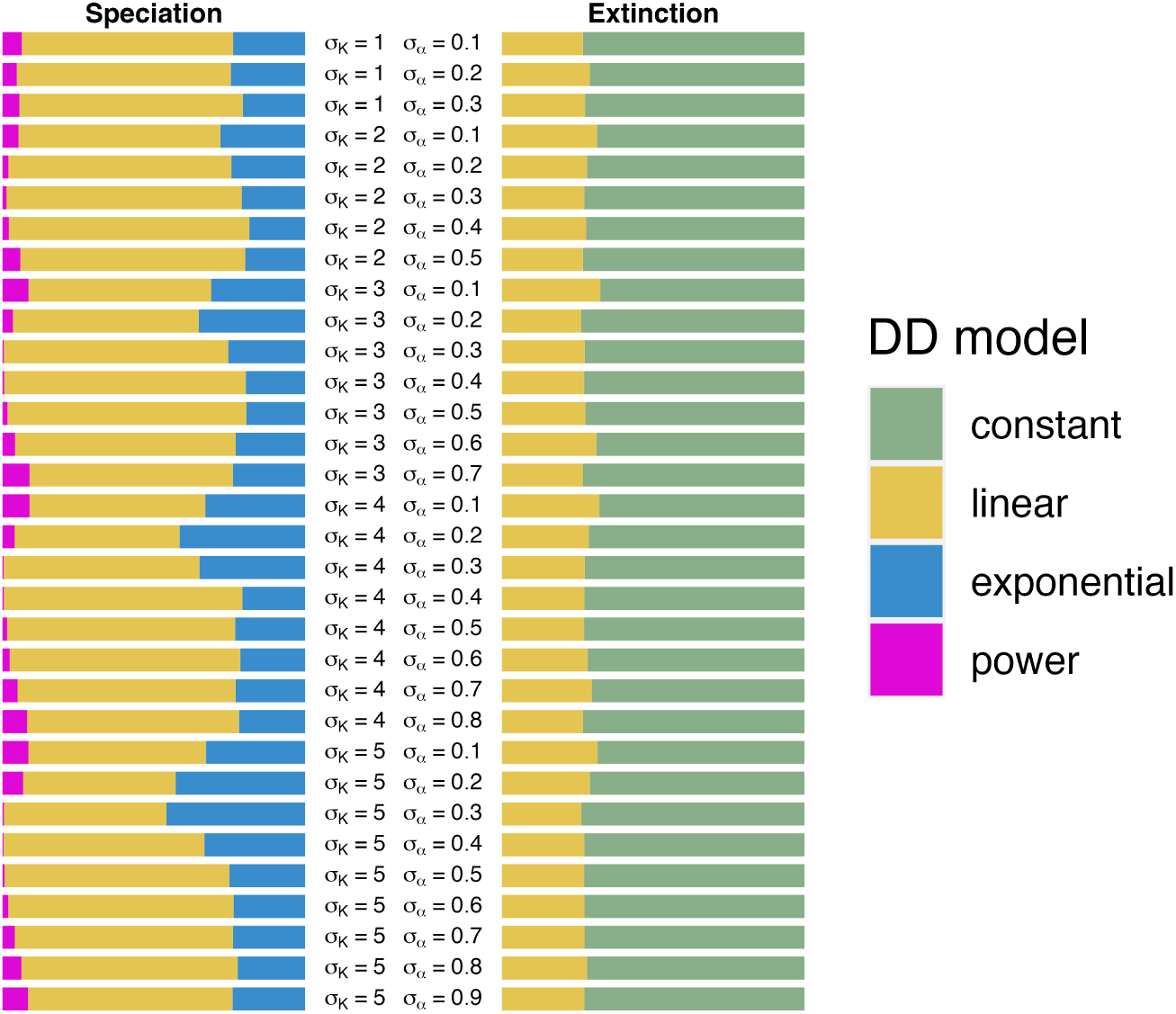
Results of model selection using birth-death models and maximum likelihood, for reconstructed trees. The width of the bars denotes the total AIC weight for each diversity-dependent function, taken as the sum of average AIC weights (across all 100 trees) of all birth-death models that included that function.

**Figure 13:**
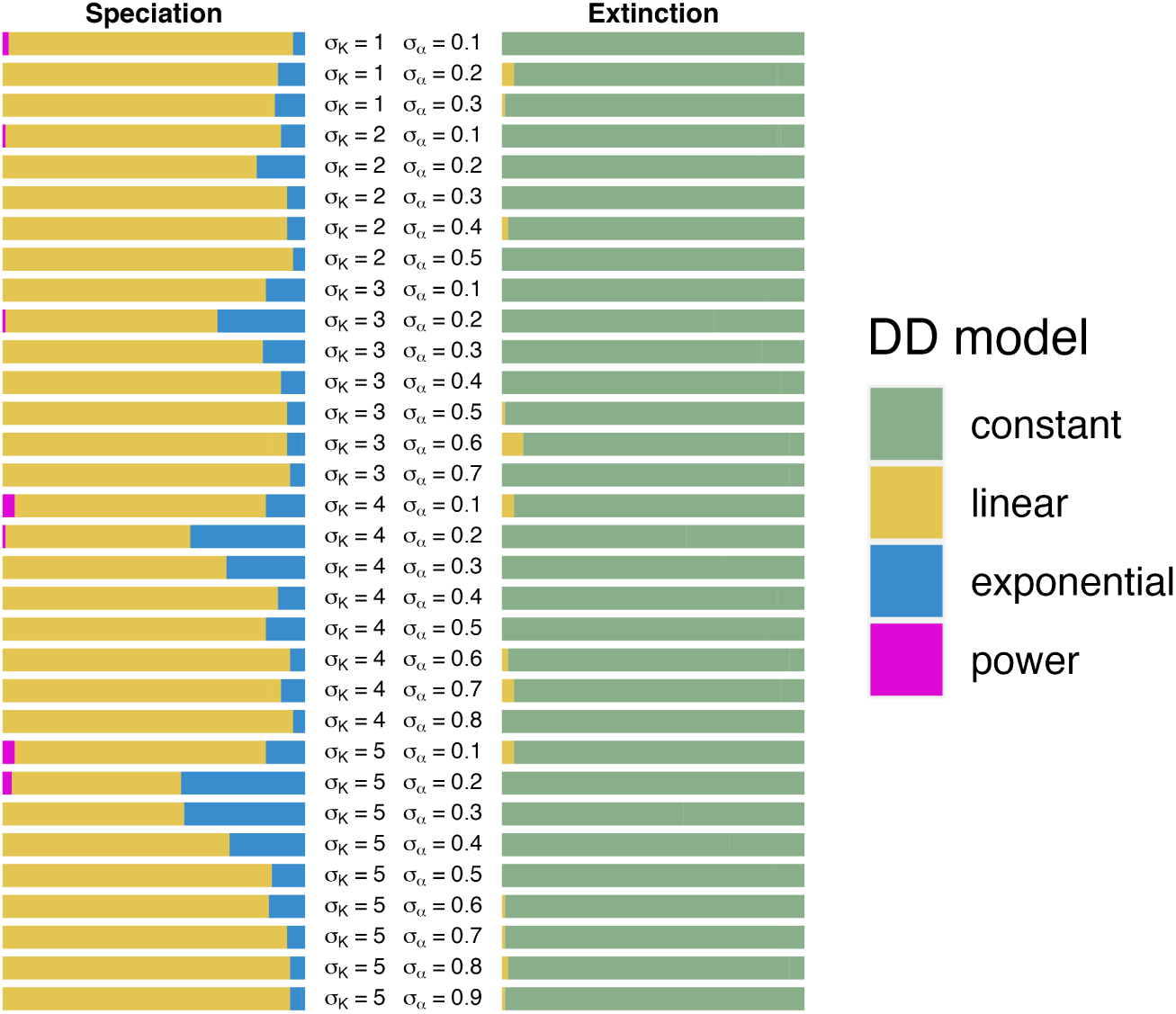
Results of model selection using birth-death models and maximum likelihood, for reconstructed trees. By contrast with 12, support for a model is measured as the total number of occurrences of this model as the best model among the 100 replicate trees.

In many cases, maximum likelihood estimates of *K* associated with power speciation models appear unreasonable, taking values over 1,000 species for many trees (Fig. 15). This concerned all parameter settings with *σ_α_ >* 0.1. The implication of these values would be the near constancy (i.e., no diversity-dependence) of the speciation rate (pink lines in Fig. 14), a conclusion that should be dismissed given the clear deceleration of speciation observed in the corresponding lineage-through-time plots (Fig. 3). Exponential speciation models display the same issue, to a lesser degree, with estimates of *K >* 100, recovering diversity-dependence but still overestimating the equilibrium diversity by a large factor (Fig. 15). By contrast, estimates of *K* associated with linear speciation models are always close to the values estimated from the complete trees and estimates of *K*^^^ (Fig. 15). This issue is however insufficient to explain the better fit of the linear model: in cases where all three speciation models yield credible estimates for *K* (that is, settings with *σ_α_* = 0.1), power and exponential diversity-dependence on speciation remain poorly supported (Fig. 12). Surprisingly, further examination of the estimated values of the parameters associated with each model does show that explosive speciation is detected by the exponential and power models in cases where reasonable values were estimated for *K* (pink and blue lines in Fig. 14). Estimated values of *λ*_0_ for reconstructed trees are indeed close to those estimated from complete trees, for all speciation functions (Fig. 15). Estimates from the linear speciation models are particularly consistent with the values estimated from complete trees. While *λ*_0_ tends to be overestimated in the case of exponential and power speciation models (up to half an order of magnitude for exponential models and up to an order of magnitude for power models), the importance of this bias is reduced by the curvature of the speciation rate specified in those models: in the end, early explosive speciation is a feature of the resulting speciation rates, and those match the early phase of the rates estimated separately for all values of *N* well. Estimates of the speciation rate in the second phase of its decline with diversity tend to decouple from the rates estimated from complete trees, largely as a result of extinction being estimated as absent (see next section). With the exception of settings with *σ_α_* = 0.1, both *µ*_0_ and *ϕ* are always estimated as close to zero (Fig. 15). In those settings and models where estimates of *K* are accurate, the consequence is that the estimated rate suggests a much steeper decline of speciation with *N*, and a lower rate of speciation at equilibrium (*N* = *K*) than what is observed in complete trees.

**Figure 14:**
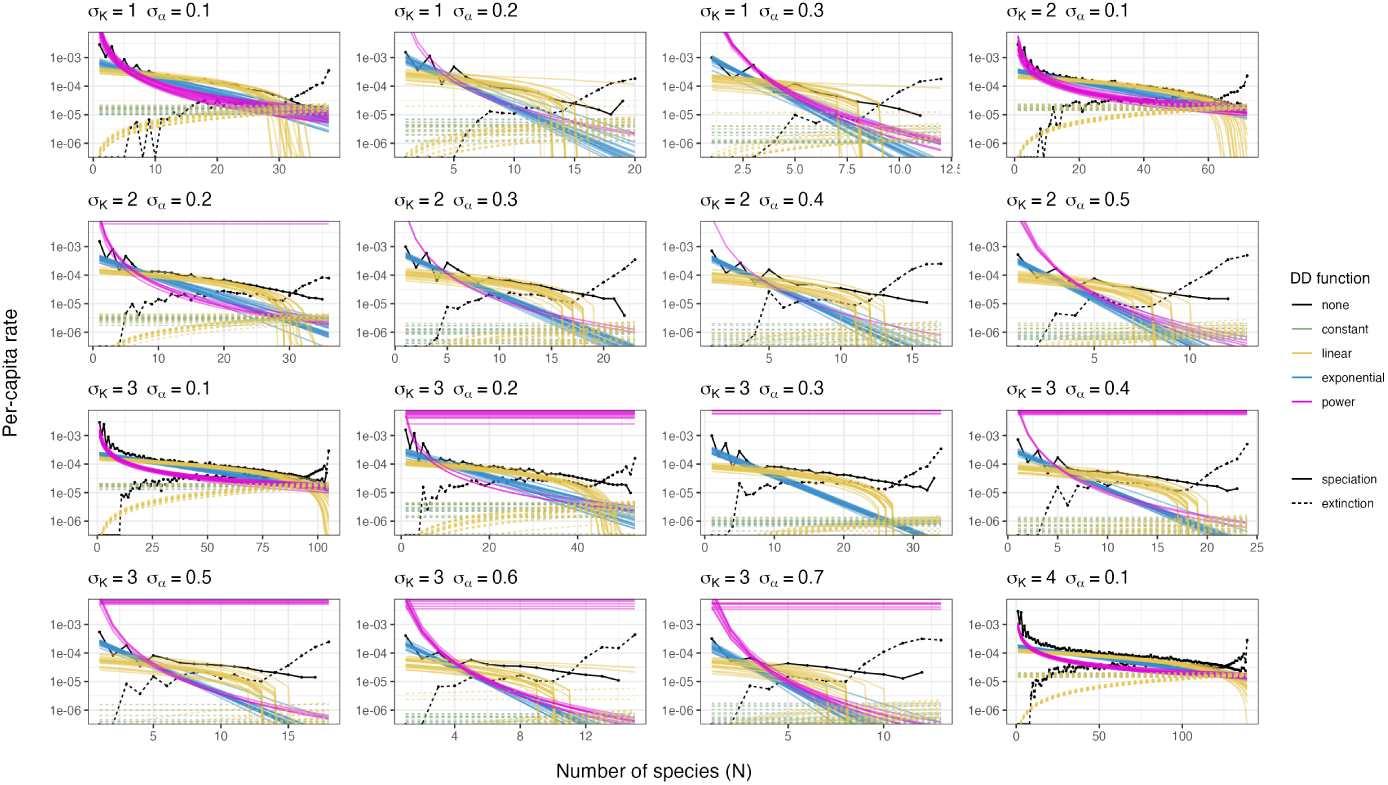
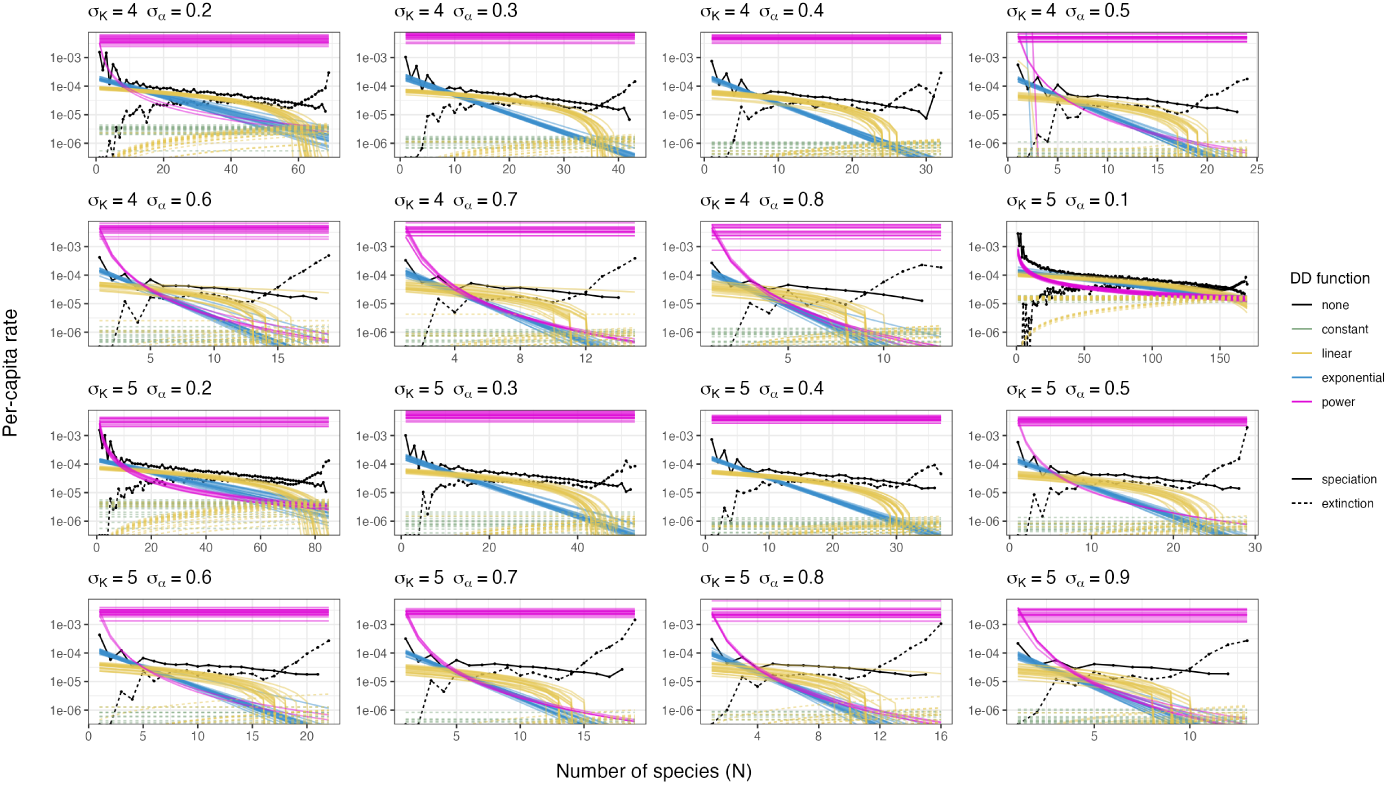
Maximum-likelihood estimates of the speciation and extinction rates obtained from reconstructed trees (coloured lines). Since models were fitted separately on every tree, plotted are the maximum-likelihood estimates corresponding to a random sample of 20 trees among the 100 replicates. Rates estimated independently for each value of *N* (Fig. 6) are plotted (black lines) for comparison.

**Figure 15:**
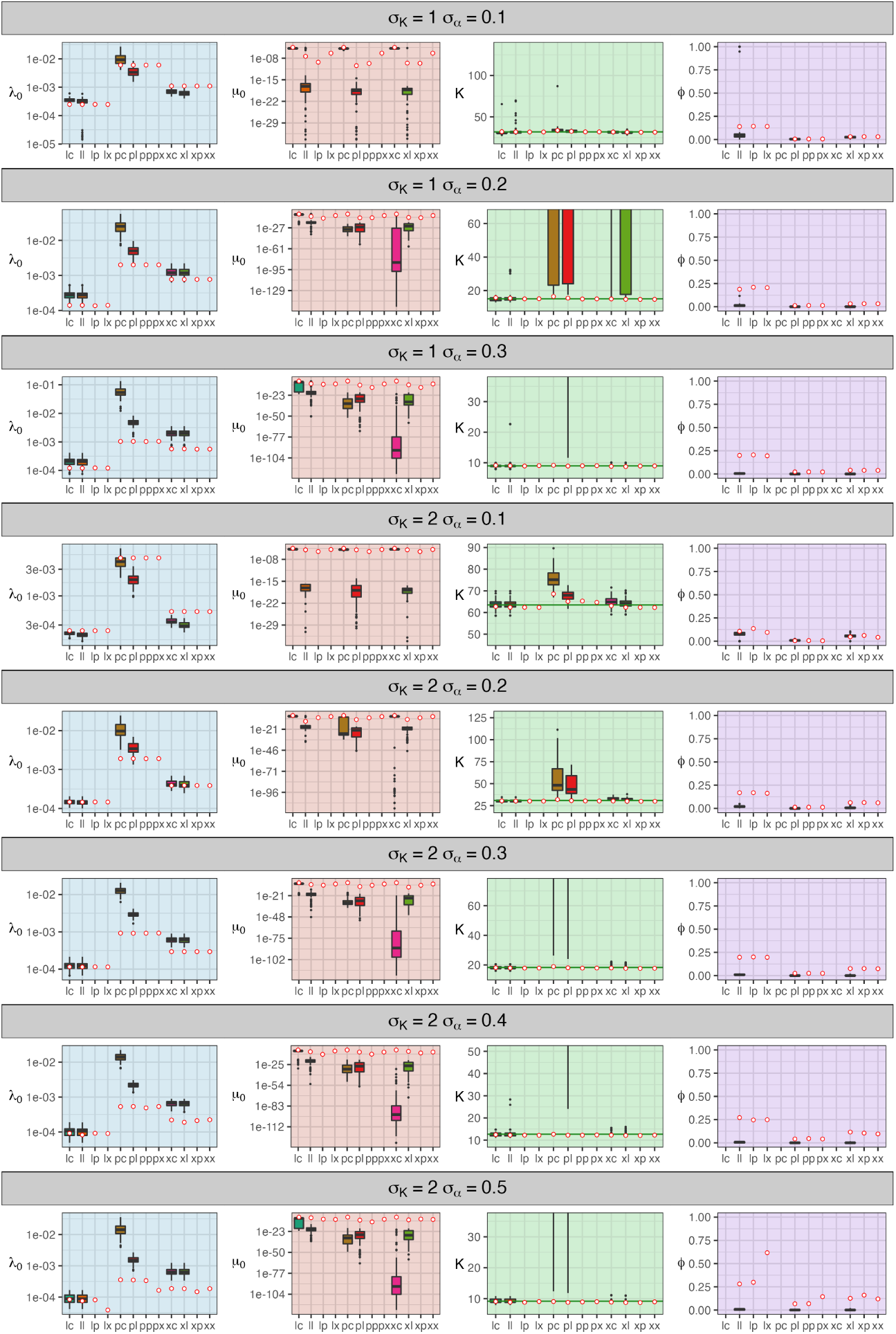
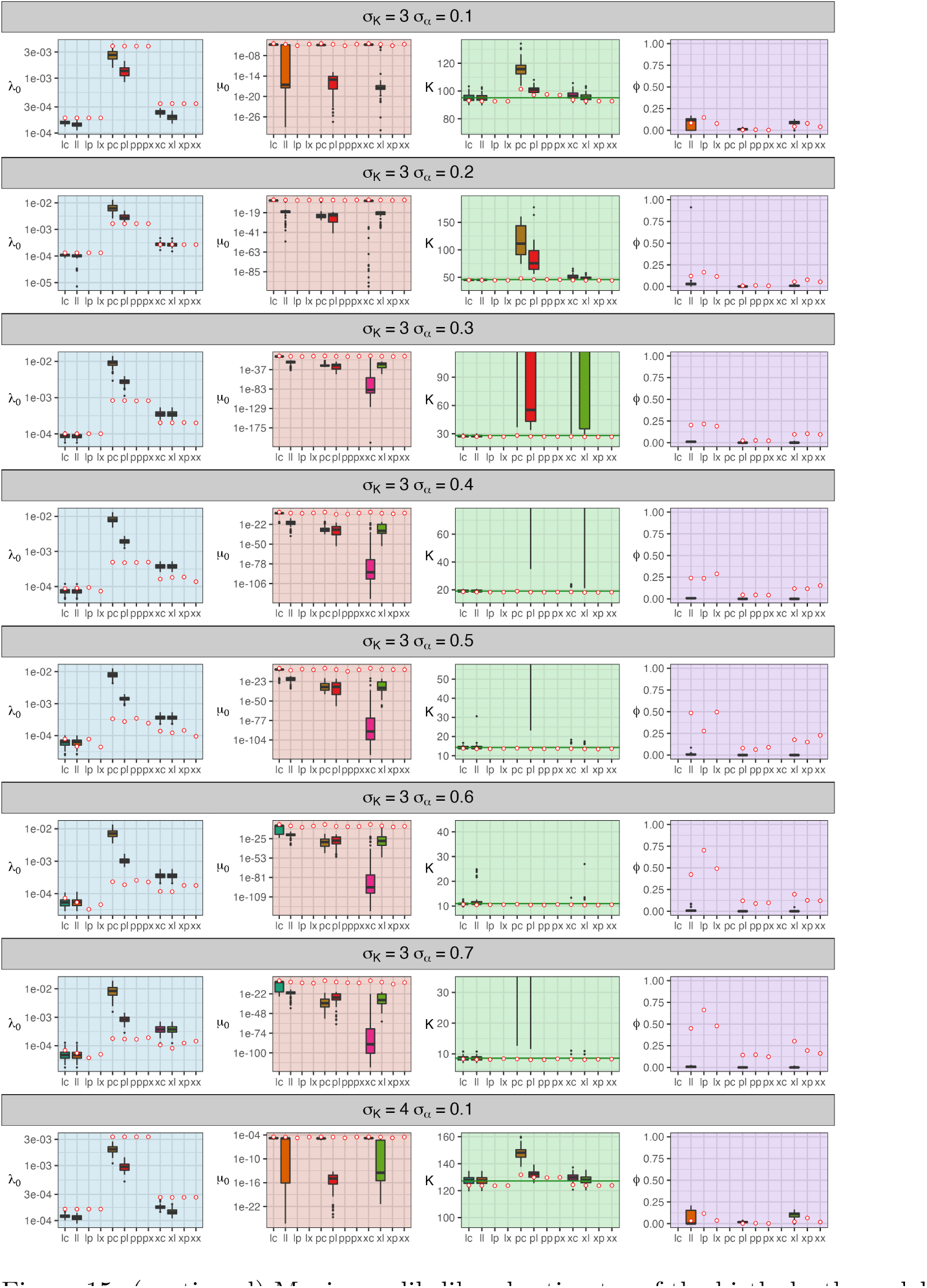
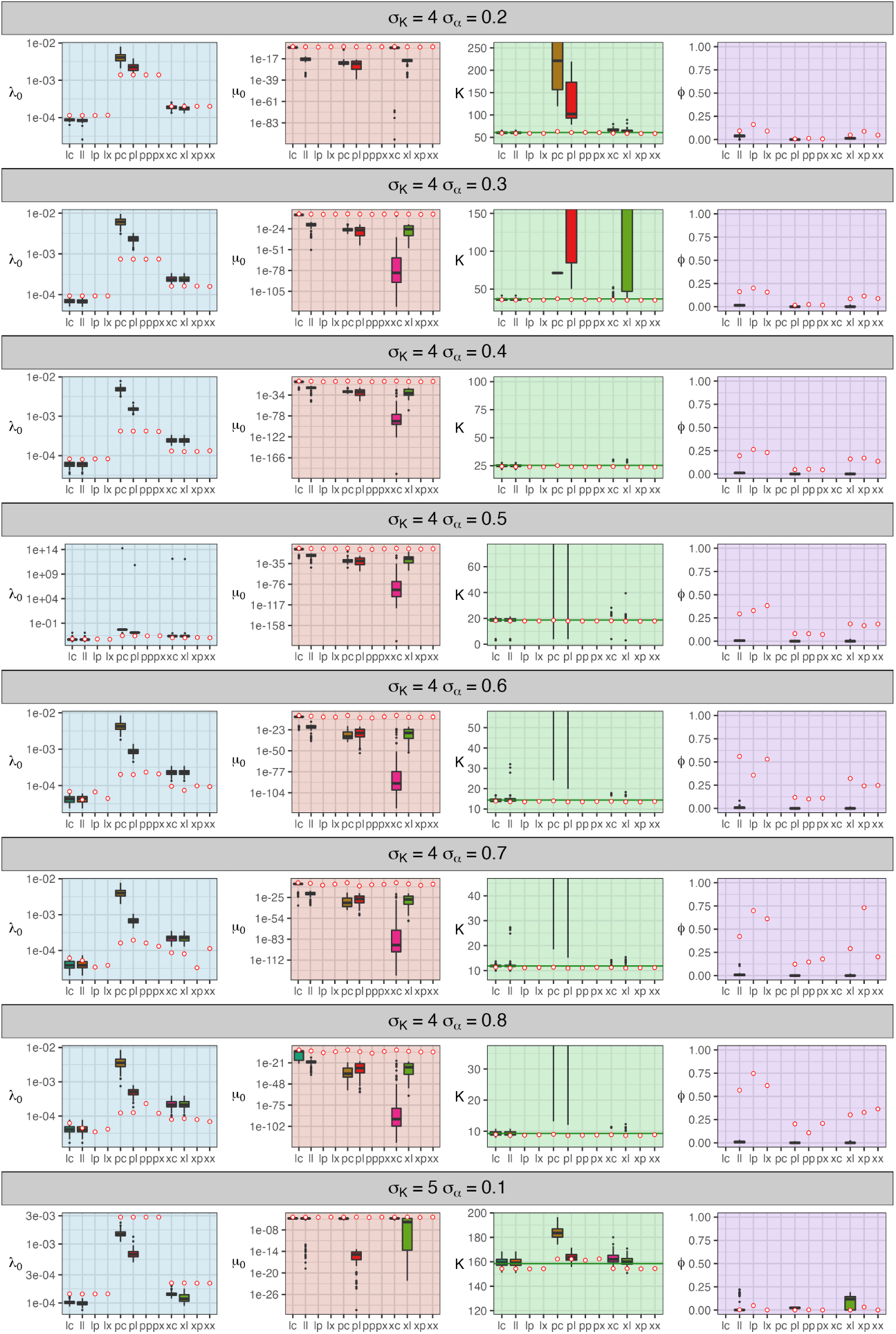
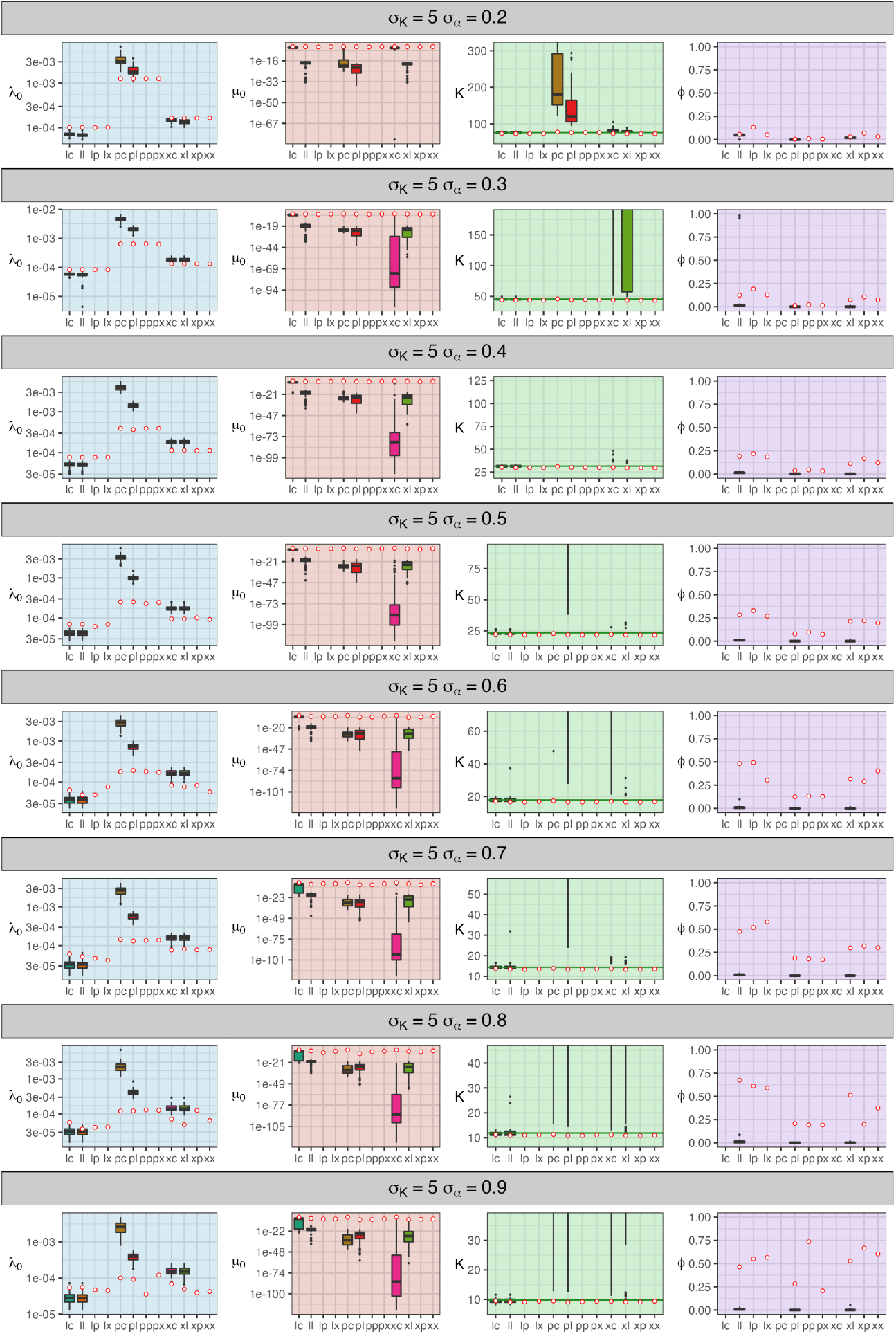
Maximum likelihood estimates of the birth-death model parameters, shown separately for each model. Models are named after the speciation and extinction function that compose them (”c” for constant, ”l” for linear, ”p” for power, ”x” for exponential). Estimates obtained from complete trees are shown as red circles (one estimate per model and set), and estimates obtained from reconstructed trees are shown as box-plots (one estimate per model and tree). The green horizontal bar in the *K* panels denotes *K*^^^ . Note that estimates for models ”lp”, ”lx”, ”px”, ”pp”, ”xp”, and ”xx” are missing for reconstructed trees as we could not obtain estimates or these models (see main text).

#### 3.4.2 Diversity-dependence in extinction is not recovered in reconstructed trees

Maximum likelihood optimization proved challenging for exponential- or power diversity dependence models. For all models that included either exponential or power diversity dependence on extinction, solving the ODE numerically became computationally intractable for some values of the parameters of the birth-death models, effectively putting the optimisation algorithm to a halt. We could not link the occurrence of this issue with any particular condition of the parameter set. We chose to exclude these models from the analysis, and proceed with a comparison of linear diversity-dependence on extinction against constant-rate extinction (right panels in Fig. 12). As a result, we were not able to infer what form of diversity-dependence on extinction would be inferred from the molecular phylogeny of a clade evolving under a scenario similar to the LVIBM, as we did for speciation above.

It can however be anticipated that the likelihood of models with the two missing forms of extinction would be close to models with linear diversity-dependence on extinction (all of them specifying that extinction should generally increase), rather than constant-rate extinction. Indeed, we observed this for complete trees (Fig.10). We thus treat the linear model as a proxy for diversity-dependence in general, and model selection becomes a test for the detection of diversity-dependence in the extinction rate against its absence.

In this perspective, we find that the diversity-dependence in extinction observed in complete trees is not recovered in reconstructed trees: constant-rate extinction is decisively supported in all parameter settings (Fig. 12). The lack of signal for diversity-dependent extinction is further confirmed by the values of parameter *ϕ* estimated for the three models with linear diversity-dependence in extinction (Fig. 15). Despite substantial variation in the estimated value of *ϕ* in the complete tree case (see previous sections), in reconstructed trees *ϕ* is almost always found to be zero (Fig. 15). This implies that diversity-dependence is carried entirely through the decline of the speciation rate, and the rate of extinction is effectively constant even in those models that assume diversity-dependence in extinction (dashed lines in Fig. 14).

Furthermore, many models did not recover extinction at all: values estimated for both parameters *ϕ* and *µ*_0_ were virtually zero (Fig. 15). This included models specifying diversity-dependent extinction, in all settings, as well as models with constant-rate extinction and either power or exponential diversity-dependence in speciation, in settings with *σ_α_ >* 0.1. Only the model with linear diversity-dependence in speciation rate and constant-rate extinction (that is, the best fitting model in most scenarios) consistently recovered non-zero extinction (”lc” in Fig. 15). Note that the selection of this model cannot be attributed to the failure of other models to infer extinction: in those settings (*σ_α_* = 0.1) where power or exponential diversity-dependent speciation models also infer non-zero extinction (”xc” and ”pc” in Fig. 15), the linear model is still strongly preferred (Fig. 12).

## 4 Discussion

Diversity-dependence in the net rate of diversification is expected to arise in an ecological scenario where resources limit diversity through increasing competition. Diversification is expected to decrease as the number of species in the community increases (McPeek 2008), but the quantitative relationship between diversity and diversification is not known. Recognizing a general decrease of diversification as a clade grows does not constitute satisfying evidence for a central role of competition in the evolution of the clade, because many unrelated, ecology- and non-ecology-based processes may also produce this pattern (Morlon 2014; Pannetier et al. 2021). Yet, only a handful of studies have attempted to measure what form of diversity-dependence would emerge from an ecological scenario featuring competition (Maurer 1989; Aguiĺee et al. 2018), and most studies seeking diversity-dependence in empirical clades assume a linear and/or power function of the number of species (though see Ezard and Purvis (2016) for more mechanistic models of diversity-dependence). In order to find out what form of diversity-dependence would be expected if competition indeed drives and limits diversification, and what simple function best approaches it, we used a simple individual-based model derived from adaptive dynamics and tracked how the rates of speciation and extinction change with the number of species in the community. In a first approach, separating the times between events and estimating the per-capita rates of speciation and extinction independently for each value of *N* allowed us to observe variation in the rates without any assumption regarding the larger trends, and lead us to make two important observations.

First, we obtained qualitative expectations for how the rates of speciation of extinction should change with the number of species in an evolutionary system where competition drives evolution. This relation features transitions between different phases rather than a straightforward, uniform function. The rate of speciation presents two phases: a first phase presenting initially explosive speciation, followed by a quick but decelerating decrease, and a second phase where speciation keeps decreasing at a slower, steady pace. After the first few branching events, the rate of extinction starts increasing quickly, but at a decelerating pace. When diversity has built up to a certain level, the rate of extinction accelerates quickly, equalling the speciation rate, which sets equilibrium diversity. The rate of extinction keeps accelerating if diversity exceeds equilibrium, and at an increasing rate as diversity while the rate of speciation keeps decreasing at the same pace. Between these two phases of acceleration of the extinction rate, extinction is either about constant or increases first and then decreases at a slow pace. Overall, the amount of change in the speciation rate as diversity accumulates greatly exceeds the amount of change in the extinction rate, such that diversity-dependence is overall, primarily carried by the deceleration of speciation. Yet, diversity-dependence in extinction is present, and not negligible. Diversity-dependence in extinction has been observed in the fossil record (Alroy 2008; Quental and Marshall 2013; Foote et al. 2018; Brayard et al. 2009), but has been poorly supported in molecular phylogenies. Here we show that competition indeed generates diversity-dependent extinction, although only for the lower and higher ranges of values of *N* . Extinction is otherwise roughly constant through a range of intermediate values of *N* . The duration of that constant-rate, intermediate phase relative to the two increasing extinction phases depends on how large is the equilibrium diversity, which in turn is determined by the ratio 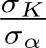. As a result, the degree of diversity-dependence in extinction depends on *α* equilibrium diversity: when it is low, extinction increases with diversity consistently, while when it is high, extinction is only diversity-dependent at low and high values of diversity.

Second, we described how the two parameters of the individual-based model contribute to modulate diversity-dependence. Surprisingly, we found that intense competition (higher values of *σ_α_*) reduced the baseline rate of speciation, independently of diversity (Fig. 6), even when the community contains a single species (*N* = 1). That is, the level of intraspecific competition directly affects the rate of divergence of populations within species. This is a counter-intuitive result, as one could expect that intense competition would increase frequency-dependence, strengthening disruptive selection. We hypothesize that this is a consequence of a smoothening effect of *σ_α_* on the selection gradient:

large values of *σ_α_* cause the frequency-dependent component of fitness to decrease more slowly away from individual clusters; resulting in a flatter fitness gradient, weaker disruptive selection, and eventually, slower speciation. However, the main effect of both parameters on the speciation and extinction rates is expressed through equilibrium diversity: higher values of *σ_K_* (increasing the abundance of resources) increase the equilibrium diversity, while higher values of *σ_α_* reduce it. More specifically, lower values of *σ_K_* and higher values of *σ_α_* increase the slope of the speciation rate in the second phase of decline; and advance the onset of the last accelerating phase of extinction, causing the two rates to intersect at lower value of *N* . An ongoing debate regarding the nature of diversity-dependence concerns whether it brings a hard limit on diversity given a finite niche space, or whether diversification is only reduced further and further as diversity rises, yet without ever stopping (Benton 2009; Harmon and Harrison 2015; Rabosky and Hurlbert 2015). The communities emerging from the LVIBM show a clear, predictable equilibrium diversity and thus support the former. Branching does not continue ad infinitum as would happen in the deterministic version of the model (Doebeli 2011). An explicit expression of the equilibrium diversity as a function of both seems to exist, as evidenced by the very low variation in the equilibrium community size across replicate simulation sharing the same values of the parameters; but we were not able not identify it entirely.

While it is clear that equilibrium diversity is a linear function of *σ_K_*, the relation between the former and *σ_α_* has a more complex form (that is, *K*^^^ = *a σ_K_ f* (*σ_α_*)). Our approximations for these functions are useful to anticipate the size of the community and choose the parameters of the model accordingly, but neither is satisfying from a biological perspective. Ultimately, an expression of the equilibrium diversity should be identifiable from the relation between the parameters and the equilibrium number of individuals in the community (i.e., the carrying capacity), which is easy to find: by dividing the area of the resource kernel (*K*(*z*)) by the area of the competition kernel (*α*(*z*)), it follows that 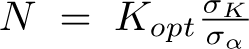. This relation was verified in our simulations, although the equilibrium number of individuals was consistently very slightly below the number predicted by this relation. At equilibrium, most species do not appear to be at a limiting population size, such that the community could in theory contain many more species, if existing populations were further divided between more species. Because the number of individuals in the community and the range of viable trait values (*z* such that *K*(*z*) ≥ 1) can both be identified, the crucial next step to identify a relation between equilibrium diversity and the parameters of the model would then be to determine how individuals are distributed in clusters (species) in the trait space and through time. In particular, we note that species are separated by gaps in trait space, putting an effective limit to the number of phenotypic clusters (i.e, species) that can coexist inside the range of viable trait values. The question may then be to identify the relation between the width of these gaps and the abundance of the species adjacent to it. While we have not succeeded in characterizing this distribution here, we hope to pursue our efforts in this direction in the future.

A number of mechanistic models of diversification have been developed in the recent years (McPeek 2008; Pontarp et al. 2012; Gascuel et al. 2015; Aristide and Morlon 2019). Here, we chose to base our model on a stochastic version of the coevolutionary models considered in adaptive dynamics studies. The choice of this model was motivated by its relevance and its simplicity: branching (i.e., diversification) occurs as a consequence of the Lotka-Volterra dynamics taking place at the level of individuals, which are controlled solely by the ratio 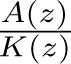. The dynamics of branching are themselves controlled by this ecological component, along with the growth rate and the rate of mutation, both of which we fixed in this study. The core of this model rests on a Lotka-Volterra coevolutionary component, that is the ratio of the competition kernel (*α*(*z*)) and niche (resource abundance) kernel (*K*(*z*)), which allows us to make connections with community- and general ecology theory. Future developments could iteratively challenge the assumptions of the model, for example switching to a sexual mode of reproduction (Dieckmann and Doebeli 1999), adding a spatial component (Pontarp et al. 2012), or explicitly modelling genetic determinism, in each case studying how each development would affect the evolutionary rates and form of diversity-dependence observed. Finally, the foundation of the model in adaptive dynamics allows us to draw predictions from existing knowledge of adaptive dynamics; for example, the equations describing the speed of evolution of the trait along a branch (and thus, the pace of divergence) are known (Doebeli 2011), and from this we can anticipate that increasing the rate of mutation (within the limit of keeping mutations “rare”, a fundamental assumption of adaptive dynamics that is a requirement for branching to occur) in the present model should accelerate evolution, and, as a result, divergence and speciation. Eventually, we hope that this work will contribute to the foundation of a mechanistic, multi-level theory of macroevolution. A call for such a theory has been made by several authors recently (Aristide and Morlon 2019; Weber et al. 2017; Hembry and Weber 2020). In such a view, IBMs such as the one presented here could be used to derive macroevolutionary predictions from ecological, contemporary models, and such predictions could be confronted to empirical patterns observed in the fossil record, the distribution of branches in phylogenetic trees and the range and trait distribution along the tips of phylogenetic trees. Eventually, a clearer view could emerge from iterative modifications of these initially simple models. This is nothing new of course, as studies aiming towards this goal have been (e.g., Maurer (1989); McPeek (2008)) and continue (Aguiĺee et al. 2018; Xu et al. 2021) to be undertaken, but a synthetic theory has yet to emerge. Arguably, the pace of branching events depends to some extent on the shape of the resource distribution function, *K*(*z*). In the present LVIBM, the resource distribution is assumed to follow a Gaussian function, a choice that is aligned with classic ecological theory. Whether or not the Gaussian function constitutes an appropriate modelling choice is a debate that exceeds the scope of this paper, but it is interesting to note that *K*(*z*) functions emerging from consumer-resource models are rarely Gaussian (Doebeli 2011). For example, multi-modal, or skewed resource distribution functions are not biologically unreasonable. This may certainly impact the pace of branching events: a multi-modal function may cause successive bursts-and-slowdowns as the clade expands into sections of trait space associated with different peaks; with a skewed distribution, branching will halt earlier on one side of the optimum, and continue longer on the other, stretching the sequence of speciation events on a longer time frame. Whether and how this is in turn affecting the form of diversity-dependence in the speciation rate, however, is unclear to us, as out of consistency with the birth-death models we have made the assumption that time does not affect the probability of speciation other than through diversity *N* (*t*), and thus we have not studied how the time sequence of events may affect diversity-dependence. The present work constitutes, to the best of our knowledge, the second attempt after Aguiĺee et al. (2018) to describe the rates of speciation and extinction, and the form of diversity-dependence that emerge from these coevolutionary models. The individual-based model by Aguiĺee et al. (2018) shares the same foundation as the one we used here (namely, the fitness function is based on the ratio 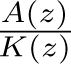), but is more complex, differing from ours in two key aspects. First, their model included a spatial component, with several sites and shifting connections between them. Second, individuals in their model reproduced sexually, and reproductive isolation was modelled explicitly. Valuable insights can be gained from a comparison of the results of our two studies. Despite of the differences between the models, we note a striking similarity between our respective results: the form of diversity-dependence reported by (Aguiĺee et al. 2018) features several phases between which the relation between diversity and the speciation and extinction rate changes. Specifically, they distinguished three phases. First, a phase of “geographic adaptive radiation”, where the initial colonization of all the sites and the resulting divergence in allopatry causes speciation to be explosive and quickly decreasing, while extinction is virtually absent. Following this, diversification enters a “niche self-structuring” phase, where local (within sites) adaptation and competition result in a quick increase of extinction, followed by about constant-rate speciation and extinction. Finally, as local niches saturate, speciation slows down, while extinction blows up, precipitating equilibrium diversity. As described above, we did find the same variations in the rates of speciation and extinction with diversity, with the exception that in our results, we did not find the speciation rate to be constant, and its slowdown phase instead appears to start immediately after the initial explosive phase. It is also possible that this discrepancy is due to a different interpretation of our respective results: we note that depending on the parameter values, constantrate speciation is not always visible in the results of Aguiĺee et al. (Fig S5 in Aguiĺee et al. (2018)). Consistent with our results, Aguiĺee et al. (2018) also reported an effect of 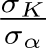 on total diversity, the speciation rate and the extinction rate (Fig. S5 in Aguiĺee et al. (2018)). Their results also show a predictable relation between 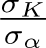 and equilibrium diversity (Fig. S4, panel h) that is at least qualitatively consistent with our results (compare with Fig. 2). Taken together, the similarities in the variations of rates between their model and ours suggest a general pattern of diversity-dependence for this class of models, and this is probably something one wants to look for as evidence for diversity-dependence. The *ad hoc* nature of this pattern we uncovered however implies it cannot be tested directly. In section 4.1 we discuss how well it is approximated by traditional phenomenological models of diversity-dependence.

We found that a significant proportion of extinction events concern recently diverged species through exclusion by their sister species, with a strong effect of the intensity of competition on the frequency of extinction of these short-lived species (the proportion of these events grow from 20% to 57% in settings with the largest values of *σ_α_*). This is consistent with the concept of “ephemeral speciation” (Rosenblum et al. 2012; Rabosky 2013) and Darwin’s own model of macroevolution (Darwin 1859; Reznick and Ricklefs 2009), which Rabosky (2013) described as a mechanism by which diversity-dependence takes place (that is, “Darwinian diversity-dependence”). Excluding ephemeral species from the data does reduce both the speciation rate and the extinction rate in equal measure, but does not change their qualitative variations with the values of *N* (Fig. 9). The different phases of speciation and extinction we have described above can still clearly be identified (Fig. 9). Therefore, while ephemeral speciation is an ubiquitous feature of the diversification process produced by the LVIBM, it does not qualitatively change the relation between the speciation or extinction rate and the number of species, so that its contribution to diversity-dependence is negligible. This finding is at odds with the results of the lineage-level mechanistic model of Aristide and Morlon (2019), where an increase in the extinction of incipient lineages was largely responsible for diversity-dependence. Arguably, the frequency of ephemeral species is sensitive to the species recognition threshold we have used. Graphically, we have observed that populations diverging from one another within species (before speciation is recognized, i.e., incipient species) are equally likely to go extinct, such that there is a continuum in the distribution of length of branches going extinct before or after speciation. Ezard et al. (2012) have shown how the choice of the criterion used to delineate species in the fossil record can affect the measured rates of speciation and extinction, including the slope of diversity-dependence. Here, justifying our species delimitation criterion is made easy by the morphological patterns produced by the LVIBM: the distribution of individuals in trait space and through time forms fairly discrete, separated clusters of individuals (Fig. 1). The only arbitrary choices we have made are to consider that species persist through speciation events as one of the resulting diverging lineages, and the speciation recognition threshold, which we chose to fix to *θ*(*z*) = 0.1 following Pontarp et al. (2012). Arguably, the latter is affecting the rates of speciation and extinction we have measured: lowering the threshold would result in the recognition of incipient species as full species, thus adding many speciation and extinction events to the data. However, while changing the value of *θ*(*z*) = 0.1 would certainly increase or decrease the base rates of speciation and extinction, it would not change the relation between the rates and *N*, and we are confident that the form of diversity-dependence we report is robust to this parameter. The ubiquity of extinction of ephemeral species was also reported by Aguiĺee et al. (2018), and thus it seems to be a feature of this class of coevolutionary models.

### 4.1 Evaluating the support for phenomenological models

The description of the form of diversity-dependence that emerges from the LVIBM we have carried out above has yielded valuable insights, suggesting some signature features of this evolutionary scenario to search for in empirical data. In the absence of an explicit expression for the rates of speciation and extinction, however, it is not possible to test support for this model in empirical data directly. Phenomenological models of diversity-dependence by contrast have been routinely used to test for diversity-dependence in phylogenies and fossil diversity are valuable to assess general trends in the data, but lack strong theoretical foundation. Since the initial propositions of Sepkoski (1978) and Maurer (1989) for the linear and power forms of the model respectively, there has been no investigation of whether these two models indeed constitute satisfying approximations of the assumed diversification mechanism. Critically, uncertainty has persisted over which version of the model is the appropriate form to represent the effect of competition: the two models have largely been used interchangeably, and often simultaneously, with identical conclusions in case either was found to better fit the tree. At times (Rabosky and Lovette 2008), quantitative differences have been mentioned (the rate of decline of speciation declining with *N* in the power model), but the potential implications have seldom been explored. In the presence of perfectly complete data, our model decisively supports either the exponential version we have introduced, or the power version, depending on the choice of parameters of the LVIBM. Support appears to result from the very high speciation rate at low diversity present in the simulations, which linear diversity-dependence does not capture satisfyingly. While speciation does decline fast with the number of species, the decline of speciation with *N* itself does not decelerate in simulations, as would be expected in the power model, and instead speciation past the initial explosive phase is best approximated by the linear model, and to a lesser extent the exponential model. Selection of exponential diversity-dependence therefore appears to be a matter of trade-off between these two features, the exponential model producing diversity-dependence intermediate between those specified in the power and linear models. Considering that none of the three types of functions capture all features of the rates estimated for each N individually, and the discrepancy between the two phases of speciation, perhaps the most appropriate birth-death model would be one including a transition between two modes of diversity-dependence. Birth-death models featuring temporal transitions between evolutionary modes have been developed (BAMM, Rabosky (2014)). Although BAMM (or any other model) does not incorporate transitions along diversity rather than time, and developing such a framework is beyond the scope of this study, comparing the fit of such a model to the ones we have included here could bring interesting insights. In the meantime, among models featuring a single mode of diversity-dependence, exponential or power diversity-dependence appears to provide the best approximation. In any case, the strong signal for explosive speciation is at odds with results reported from reconstructed phylogenies where, as we discussed in the introduction, linear diversity-dependence in the speciation rate is most often found to fit the phylogeny best. This discrepancy between our results for complete trees and empirical findings can be resolved by considering that reconstructed trees are only a limited subset of the evolutionary history of the clade. Indeed, when we perform the model selection procedure again after pruning extinct lineages from the trees, we find decisive support for linear diversity-dependence on speciation. This implies that while the signal for early explosive speciation is present, support for it is lost.

Arguably, the completeness of our assessment of phenomenological models for reconstructed trees was restricted by the two unresolved computational issues mentioned in the Results section. Below we expand a bit further on these issues, including what we suspect caused them and potential tracks to address them. First, for most trees and parameter settings, maximum likelihood estimates of the equilibrium diversity associated with exponential or power diversity-dependence on speciation are extremely high, and functionally equivalent to infinity, indicating no diversity-dependence on speciation. Paradoxically, values of parameter *ϕ* associated with this are also close to zero, indicating no diversity-dependence on extinction either. This apparent paradox can be resolved by considering the asymptotic behaviour of the speciation rate under these models. As shown by the values of maximum-likelihood estimates of the initial extinction rate *µ*_0_, the signal for weak, or even absent extinction from the reconstructed trees is strong. As a result of the formulation of the models (see Section 2.6), low values of *µ*_0_ are only possible if *K* is very high (in fact, *µ*_0_ = 0 is only possible if *K* is infinite). A potential solution to this would be to alter the diversity-dependent model and use parameters that are not susceptible to this behaviour. For example, one could define *K* as the value of *N* for which the speciation is a certain fraction of the initial speciation rate *λ*_0_, thus circumventing the issue by decoupling speciation from extinction. Such a model could be helpful to quantify diversity-dependence, but unfortunately loses the biological interpretation of the equilibrium diversity.

Second, the likelihood could often not be computed for models that incorporated exponential or power diversity-dependence on extinction, and we excluded these 6 models from the initial set of candidate models. The issue appears to originate from the extinction rate growing very large under some values of the parameters explored during optimisation. High extinction requires keeping track of the probabilities of a larger set of possible states, such that the size of the system of equations grows and integration of the likelihood becomes computationally challenging. An evident solution would be to attempt to limit the size of the system of equations to a maximum value, and forego the computation of probabilities associated with high (and unlikely) values of N. Because this will inevitably bias the likelihood to some extent (as we do not calculate part of the probability density function at every step), making it difficult to apply in practice. Nevertheless, the results we do have at hand suggest that our conclusions are robust to these missing results. In the few cases where we did obtain reasonable estimates of the carrying capacity for models that incorporated exponential or power diversity-dependence on speciation, linear diversity-dependence was still largely preferred. Constant-rate extinction was largely preferred over linear diversity-dependence on extinction, such that it appears unlikely that any other form of diversity-dependence on extinction would fit the data better.

